# A population code for semantics in human hippocampus

**DOI:** 10.1101/2025.02.21.639601

**Authors:** Melissa Franch, Elizabeth A. Mickiewicz, James L. Belanger, Brad Joiner, Kalman A. Katlowitz, Hanlin Zhu, Ana G. Chavez, Assia Chericoni, Danika Paulo, Eleonora Bartoli, Suzanne Kemmer, Steven T. Piantadosi, Nicole R. Provenza, Jay A. Hennig, Benjamin Y. Hayden, Sameer A. Sheth

## Abstract

As we listen to speech, our brains track the meanings of the words we hear.

Recent successes of large language models suggest that distributed population geometry can capture rich semantic relationships between words. Motivated by this idea, we hypothesized that semantic information in the brain may likewise be expressed in distributed patterns of activity across neurons, rather than in the activity of neurons narrowly tuned to a specific word. We recorded responses of hundreds of neurons in the human hippocampus while participants listened to narrative speech. We find encoding of contextual word meaning in the simultaneous activity of neurons whose individual selectivities span multiple unrelated semantic categories. Decoding and population geometry analyses revealed distinct neural coding principles for low-versus high-frequency words, likely reflecting the greater polysemy of common words. Similar to embedding vectors in semantic language models, distance between neural population responses correlates with semantic distance; however, this effect was only observed in contextual embedding models (GPT-2 and BERT), suggesting that the semantic distance effect depends critically on contextualization. Consistent with this, we find that neural population activity supports a multidimensional semantic subspace that aligns most closely with the contextual structure captured by GPT-2. Moreover, for semantically similar words, even contextual embedders showed an inverse correlation between semantic and neural distances; we attribute this pattern to the noise-mitigating benefits of contrastive coding. Ultimately, these results provide a neurocomputational account for understanding how neural populations track word meaning.

## INTRODUCTION

The principles by which neurons encode meaning remain poorly understood. One hypothesis is that the brain uses a population code, so that word meanings are represented in patterns of activation across a large number of units (Frisby et al., 2023; Goldstein et al., 2022, 2024; Landauer & Dumais, 1997; Mitchell & Lapata, 2008; Piantadosi et al., 2024; Pulvermüller, 1999; Rogers & Mcclelland, 2004). Interest in population coding has grown in recent years because it overlaps with how meaning is represented in large language models (LLMs, Devlin et al., 2018; Joulin et al., 2016; Mikolov, Chen, et al., 2013; Mikolov et al., 2013; Pennington et al., 2014; Radford et al., 2019; Vaswani et al., 2017). Population coding facilitates solutions to several important problems including similarity judgments, analogies, generalization, hierarchical reasoning, pattern completion, and efficient learning (Chen et al., 2017; Kommers et al., 2015; Mejri et al., 2024; Tversky, 1977; Piantadosi et al., 2024). For example, if population codes are structured so that words with more similar meanings evoke more similar neural responses, then judging similarity can be as simple as calculating the distance between two words’ response vectors (Devlin et al., 2018; Erk, 2012; Grand et al., 2022; Joulin et al., 2016; Pennington et al., 2014; Radford et al., 2019; Tversky, 1977; Vaswani et al., 2017).

Population coding has attracted growing interest from scholars in recent years (Ebitz and Hayden 2021; Saxena and Cunningham 2019; Rigotti et al., 2013; Fusi et al., 2016; Tye et al., 2024). Here, we hypothesized specifically that (1) neural responses collectively instantiate a representation of word meaning that is reliable across repeated contexts, but (2) varies as the context of the word changes its specific meaning.

Moreover, we hypothesized that (3) a large number (i.e., more than 1%) of neurons are involved in encoding any given concept, and (4) the same population of neurons is involved in a range of distinct concepts.

One difficulty in assessing the population coding hypothesis with linguistic stimuli is that language encoding is highly contextual (Devlin et al., 2018; Peters et al., 2018; Radford et al., 2019; Vaswani et al., 2017) and neuroscience research has only recently begun relating neural and contextual word embeddings (Chang et al., 2008; Deniz et al., 2023,Goldstein et al., 2022, 2023, 2024; Hong et al., 2024; Hosseini et al., 2024; Kumar et al., 2024; LeBel et al., 2021; Tang et al., 2023; Tuckute et al., 2024; Zada et al., 2024). Moreover, only one existing study has examined contextual embeddings at the single neuron level (Katlowitz et al., 2025). Indeed, word meanings in normal language are highly contextually dependent. For example, the word “sharp” may mean something entirely different in the phrases “sharp knife” and “sharp mind”. Contextual semantic embedders capture these nuances of language (Linzen & Baroni, 2025; Manning et al., 2020; Pavlick, 2025; Vaswani et al., 2017; Radford et al., 2019). We hypothesized that incorporating context would improve fit quality to neural representations of semantics.

We were especially interested in the hippocampus because of its well-established role in semantic and conceptual coding in humans (Duff and Brown-Schmidt 2012, Piai et al., 2016, Blank et al., 2016; Davachi, 2006; Dijksterhuis et al., 2024; Fletcher et al., 1997; Quian Quiroga et al., 2005 and 2009; Rey et al., 2025; Squire & Zola-Morgan, 1991; Tulving, 2002; Gelbard-Sagiv et al., 2008; Kolibius et al., 2023; Rutishauser et al., 2021; Courellis et al., 2024; Cohn-Sheehy et al., 2021; Binder et al., 2009; Suthana et al., 2015; Karkowski et al., 2025; Baldassano et al., 2017). Classic neuropsychological studies demonstrated that hippocampal integrity is essential for forming and retrieving semantic knowledge and relational memory (Squire & Zola-Morgan, 1991; Tulving, 2002; Davachi, 2006; Fletcher et al., 1997), and early lesion and imaging work showed its involvement in constructing relational and conceptual meaning from experience (Fletcher et al., 1997; Duff and Brown-Schmidt, 2012). Neuroimaging studies find language specific responses in the hippocampus (Blank et al., 2016), and implicate the broader medial temporal lobe in abstract semantic processing (Binder et al., 2009).

Additionally, intracranial recordings show that hippocampal activity tracks conceptual relationships during speech and language comprehension (Piai et al., 2016).

Electrophysiological recordings from single hippocampal neurons reveal they encode abstract, semantic-level information: they respond selectively to concepts rather than sensory details and generalize across exemplars (Quian Quiroga et al., 2005; 2009; Karkowski et al., 2025; Rutishauser 2021; Gelbard-Sagiv et al., 2008; Courellis et al., 2024; Dijksterhuis et al., 2024; Rey et al., 2025). This form of conceptual coding may support the retrieval of semantic associations and facilitate content-guided recall, as demonstrated across several studies (Dijksterhuis et al., 2024; Kolibius et al., 2023; Rutishauser 2021; Gelbard-Sagiv et al., 2008; Suthana et al., 2015). Building on this, recent work has shown that hippocampal populations also track higher-order narrative structure and evolving semantic content during naturalistic experiences (Cohn-Sheehy et al., 2021; Baldassano et al., 2017). Together, these findings converge to establish the hippocampus as a central node for representing semantic content, relational structure, and conceptual knowledge in service of language and memory.

We recorded responses of neurons in the hippocampus of epilepsy patients undergoing intracranial monitoring for seizure activity while those patients listened to continuous, narrative speech. We found that individual neurons have complex, multi-word tunings, and that lexical concepts are encoded by large and heteromorphic populations of hippocampal neurons. Moreover, we find that semantic embedding distance correlates with neural embedding distance, but only for language models that involve a high degree of contextual embedding (i.e, GPT-2 and BERT). These contextual language models predict word level neural population activity patterns better than static word embedding models. Indeed, neural embedding variance correlates with polysemy, changing activity with different instances of the same word. These contextually driven neural representations may contribute to the separation of high-and low-frequency words observed in population geometry and decoding models. Finally, we find evidence for a strong contrastive code for highly similar words. Together, these results endorse the hypothesis that the hippocampus employs population coding principles for semantics during natural language comprehension. In this code, semantic relationships are reflected in the geometry of neural population activity: words with similar meanings evoke more similar patterns of activity across neurons, whereas semantically distinct words evoke more divergent patterns. Importantly, these relationships align most strongly with contextual language embeddings, indicating that the hippocampal population code reflects context-dependent meanings rather than fixed lexical categories. Thus, contextualized semantic structure is recapitulated in distributed neural responses in the hippocampus.

## RESULTS

### Neural encoding of meaning

Ten native English-speaking patients (six males and four females) undergoing neural monitoring for epilepsy listened to 47 minutes of English speech (six monologues taken from the Moth podcast, **Methods, Fig. 1a and b**). This stimulus set consisted of 7,346 words, of which 1,351 were unique. We collected responses of isolated single neurons in hippocampus (HPC, n=356 neurons, **Extended Data Fig. 1**) from 10 patients. Overall, neurons showed a clear response to the onset of individual words with a characteristic ramping to a peak, followed by a gradual decline in firing, although there was a great deal of diversity across neurons (**Fig. 1c-e and Extended Data Fig. 2**).

**Fig. 1.**
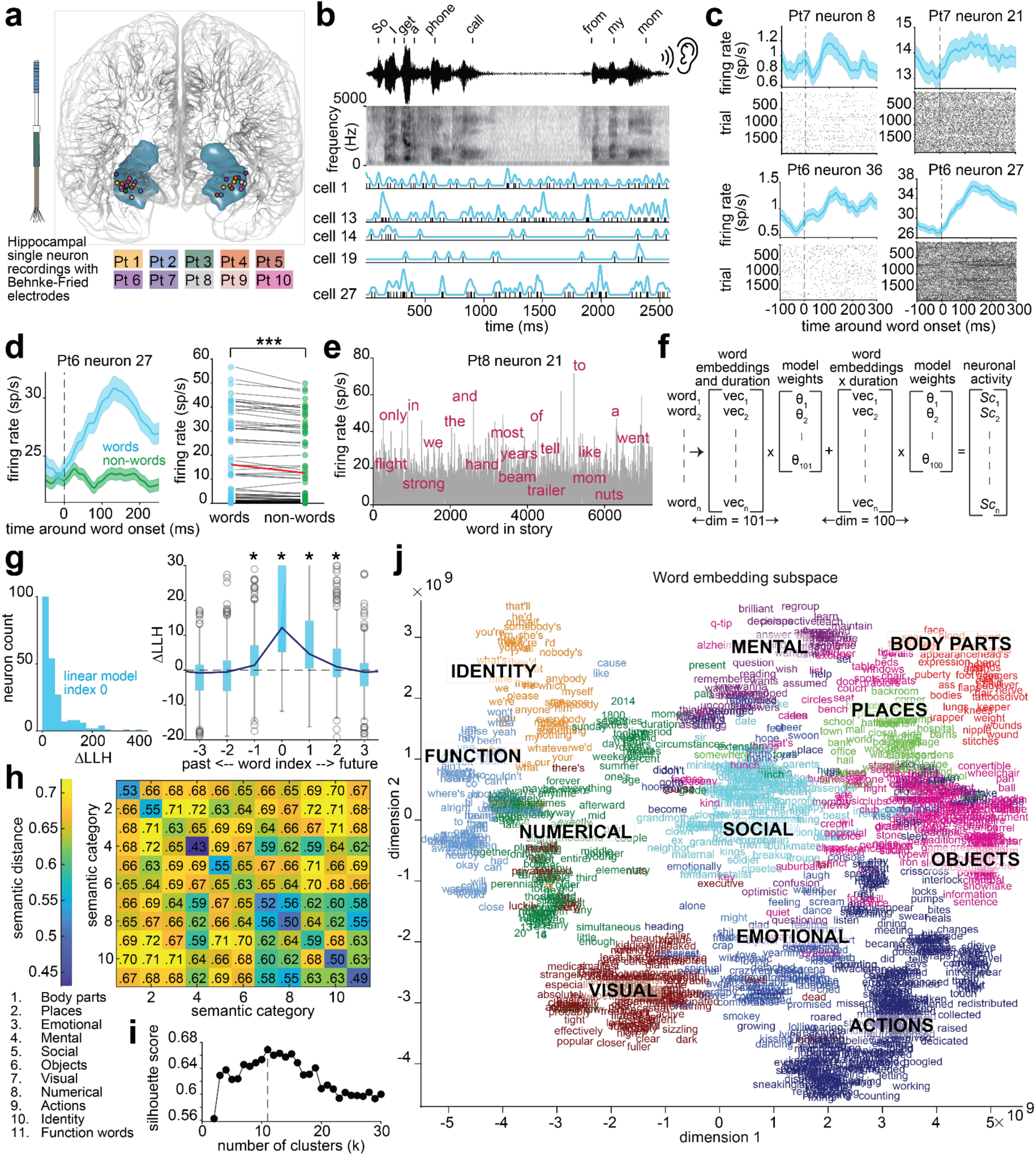
Hippocampal neurons track spoken words during language comprehension. a, Electrode recording location of hippocampal neurons from ten patients. **b,** Continuous spiking activity from five example hippocampal neurons from patient 6 during words heard during the experiment, as shown in the spectrogram. **c**, Peri-stimulus time histograms and raster plots demonstrating responses to word onsets from four distinct hippocampal neurons in patient 6 (bottom row) and 7 (top row). Dashed lines represent word onset. Plots display mean firing rate ±SEM shaded. **d,** Left - Peri-stimulus time histogram during words and jabberwocky (nonsense) non-words. Right - 71/125 neurons from four patients that exhibit significantly different firing rates between real and non-words. P = 5.6×10^-5^, Wilcoxon signed-rank test. Red line represents the mean firing rate from words and non-words (14.78 and 13.98). **e**, The firing rate of one neuron occurring across the duration of all words from the six stories during language comprehension. **f**, Regression equation used to fit an individual neuron’s spike count from word embeddings, word duration, and their pairwise interactions. Dim reflects the number of predictors including the first 100 principal components of the word embeddings and word duration. *SC* is a neuron’s spike count for each word. **g**, Left – histogram of log-likelihood improvements (LLH diff; actual – shuffled models) for fitted neurons (n = 222) from Poisson ridge regression predicting neuronal spike counts using the equation in **f** for the current word. Right – Log-likelihood improvement of all neurons from the current word duration and embeddings (index 0) and embeddings of past and future words. Boxes reflect the interquartile range with median; navy curve is fit to the median of each box; circles are outliers. The y-axis is truncated for visual clarity. **h**, Similarity matrix of median pairwise cosine distances between word embeddings within and between semantic categories that arise from all words. **i**, Silhouette score used to determine the number of semantic categories for K-means clustering of words. **j,** All words plotted in their dimensionality reduced embedding space and colored according to their semantic category.

**Fig. 2.**
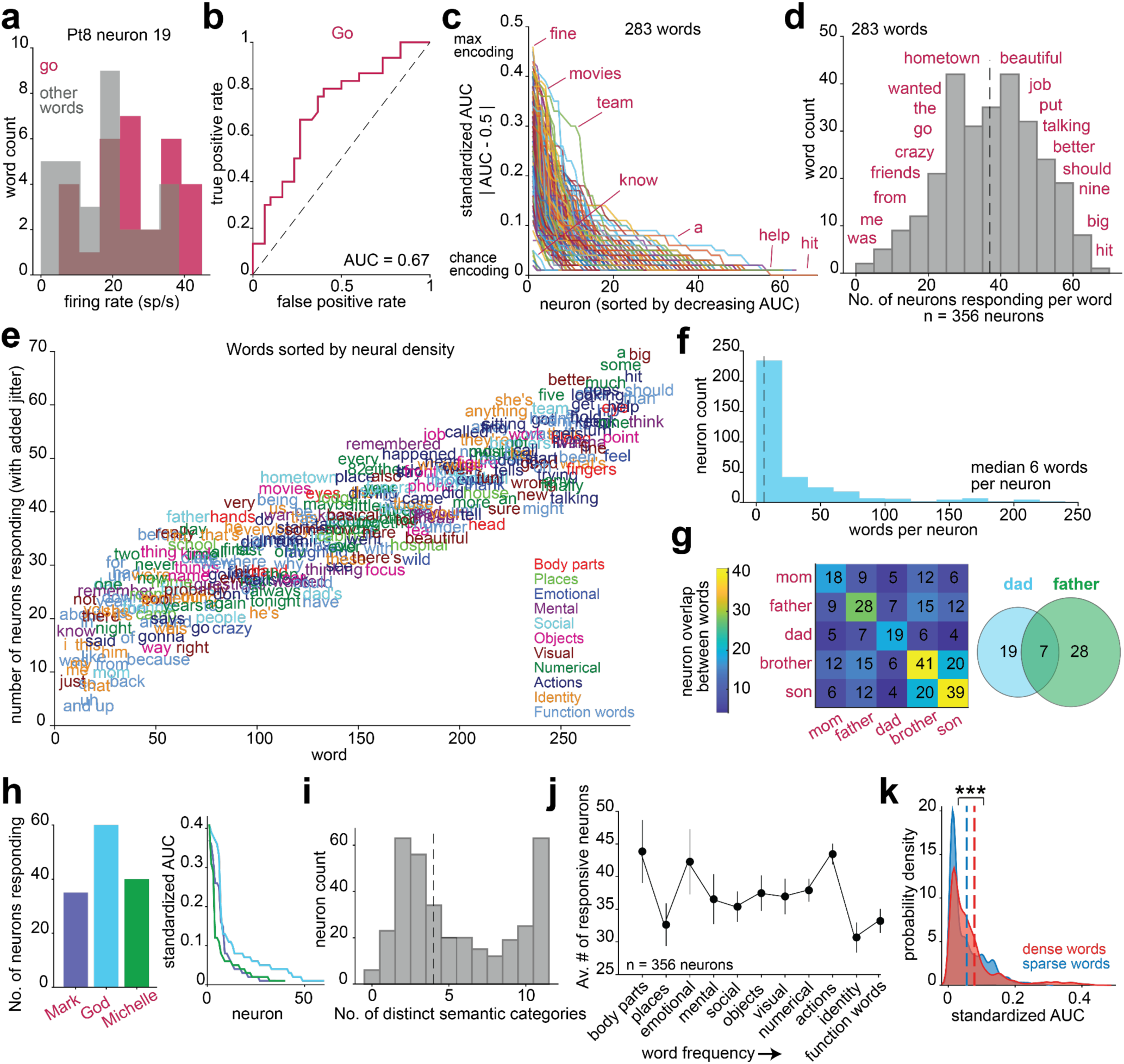
Hippocampal neurons represent words with a distributed code. **a**, Example firing rates from one neuron during all occurrences of the word, “go” and a random subset of other words. **b**, Receiver operator characteristic curve for the same neuronal activity and words in **a**. **c**, For each word (line), the standardized area under the curve (AUC) from each neuron that significantly classified that word from all others (see Methods). For each word, neurons encoding that word are sorted on the x-axis from greatest to lowest encoding performance. **d**, Histogram of the number of neurons encoding each word. Every word was encoded by at least 3 neurons. **e**, The same as in **d** but all 283 word text plotted based on the number of neurons that encode them, with added jitter for readability. **f,** Histogram of the number of words encoded by each neuron. **g,** Assessment of overlap between neurons encoding words related to the concept, family. Numbers in the matrix and circles represent the number of coding neurons. **h,** Left - Number of neurons encoding proper nouns from ROC analysis. Right - AUC values for proper noun encoding. **i,** Number of semantic categories from the words that each neuron encodes. Dashed line represents the median. **j,** Average number of neurons encoding words from each semantic category. **k,** Probability density plots of AUC values from sparse words (n = 1,517 AUC values from words with < 28 cells encoding them) and dense words (n = 1,632 AUC values from words with more than 55 cells encoding them). P = 1.643×10^-9^, Wilcoxon rank-sum test.

Across words of different durations (short ∼100 ms, middle ∼200 ms, long ∼500 ms), neurons show a similar rise in firing rate shortly after word onset, but responses attenuate quickly for short words and remain sustained for longer words (**Extended Data Fig. 3**). Therefore, we computed firing rate during an epoch starting 80 ms after the onset of each word (to account for the approximate response latency) and lasting the duration of the word (see **Methods**). Rates were normalized by word duration. All analyses were repeated with a variety of response epochs which yielded similar results (data not shown).

A subset of patients (n=4) also listened to monologues containing nonsense words (Keuleers & Brysbaert, 2010, *jabberwocky stimuli*, 1,890 words, see **Methods**) following the podcast episodes. The nonsense narratives included words modified from real ones to create meaningless, novel terms, such as “toom” or “skook.” These nonword stimuli preserved syntactic structure by retaining function words and morphemes but lacked coherent semantics. We found that 57% of neurons (75/125) exhibited a significant change in firing rate to non-word stimuli, with most cells showing markedly reduced activity for nonwords relative to words (**Fig. 1d and Extended Data Fig. 2b**).

Additionally, hippocampal neural population activity strongly distinguishes words from nonwords, achieving an average decoding accuracy of 75% (chance is 50%; **Extended Data Fig. 2c;** see **Methods**). Note that these Jabberwocky stimuli do not fully control for language because listeners could have stopped paying attention to them once they realized the stimuli did not carry meaning.

Next, we sought to better understand the relationship between neural activity and semantics, particularly how language unfolds over time as words are heard in sequence. To do this, we predicted each neuron’s spike count for a given word from its dimensionality-reduced Word2Vec embeddings (Mikolov, et al., 2013), word duration, and their interactions (**Fig. 1f**). Word2Vec represents each word as a fixed (i.e., non-contextual) 300-dimensional vector, meaning, for example, the embedding for “camp” remains the same whether it is used as a noun or a verb. The Poisson linear model (cross-validated, ridge regression; mixed effects models are not used since we fit a model to each neuron from each patient separately) fit the responses of 62% (n=222/356) of neurons, showing a significant and positive log-likelihood difference between actual and shuffled models and positive adjusted R-squared values (**Fig. 1f-g; Extended Data Fig. 4c** see **Methods**). Semantic encoding models explained a median 26% of the neural variance (**Extended Data Fig. 4c**). Adding semantic predictors to a duration-only model significantly improved fit, raising adjusted R² by a median of 0.01 (Wilcoxon z = 11.35, p < 10⁻⁹) and improving model performance by an average of 23% across neurons (**Extended Data Fig. 4d**). Likewise, semantic information significantly increased the Pearson correlation between actual and predicted spiking activity by 0.02 (semantic model median r = 0.68; Wilcoxon signed-rank z = 5.99, p < 10⁻⁹). Together, these effects indicate that semantics captures variance in neural responses that cannot be explained by word duration alone. We found comparable results with contextual embeddings (e.g., BERT embeddings fit 62% of neurons and GPT-2 layer 37 fit 54% of neurons; Devlin et al., 2018; Note that we extract BERT embeddings in a left-to-right, unidirectional manner for all analyses in this study, as seen in Lyu et al., 2023, Madureira et al., 2024 and Kahardipraja et al., 2021.). Across patients, the model fit an average of 60% of recorded neurons, with all patients contributing fitted neurons (mean = 10% recorded neurons per patient). Among neurons successfully fit by the model, significant predictors included word duration, embedding dimensions, and their interactions. Notably, the first eight embedding dimensions were commonly selected across neurons, with the number of significant predictors gradually decreasing across higher dimensions (see **Extended Data Fig. 4a-b and Methods**). Removing function words and pronouns produced a modest but significant drop in model fit (**Extended Data Fig. 5a–b**), yet their exclusion did not alter any main findings (neither here nor throughout other analyses in the paper; we therefore retain them).

To investigate whether neurons encoded the semantics or phonetics of a word, we also fit neural tuning models with the phonemes of each word as predictors instead of their embedding values (see **Methods**). Phonetic models fit fewer neurons than semantic, fitting 42% (n=150/356) of neurons. Additionally, for neurons fit by both models (40%, n = 142/356), the semantic model had significantly higher performance than the phonetic model (median phonetic – semantic ΔAIC = 136, p < 0.001 and **Extended Data Fig. 4e-f**). We find that, across neurons, adding semantic predictors significantly improved model fit relative to a phonetic-only model (median ΔR² = 0.003, sign-rank z = 12.4, p < 0.001). In contrast, adding phonetic predictors to a semantic model did not improve performance and slightly but significantly reduced explained variance (median ΔR² = –0.001, z = –2.7, p = 0.006; **Extended Data Fig. 4f**). Together, these results indicate that neural responses are better explained by semantic than phonetic features, suggesting that single-neuron activity reflects the meaning rather than the sound structure of words.

To understand the relationship between hippocampal activity and semantic, phonetic, and syntactic features, we applied the same ridge regression encoding models with a total of 61 linguistic features. Specifically, models included (1) full phonetic embeddings (43 IPA phones; count coded, cf. Khanna et al., 2024), (2) 10 PCs of semantic embeddings (Word2Vec, BERT, or GPT-2), (3) word order in sentence, (4) word order in clause, (5) opening node number, (6) closing node number, (7) word frequency (SUBTLEX-US), (8) pitch, (9) a scalar representing degree of polysemy (Cruse et al., 1986; Landauer, 2001; Ethayarajh, 2019), and (10) word duration (Manning et al., 2014). We found that one or more semantic embeddings were significant predictors for a mean 64% of the neurons fit by the model, followed by phonetic embeddings as the next most common significant predictor across neurons, fitting 41% of neurons on average (**Extended Data Fig. 4g**). Semantic embeddings also had the highest weights, followed by phonetic embeddings and opening and closing nodes (**Extended Data Fig. 4h)**. Linguistic features like pitch, word position in clause, and word frequency were significant predictors for only 18% of neurons on average and had lower weights than semantic and phonetic predictors. Moreover, after regressing out variance explained by non-semantic linguistic features and modeling the residual neural activity, semantic embeddings continued to significantly predict firing (92/356 neurons; adjusted R² = 0.08; r = 0.35). In summary, even with a variety of linguistic features accounted for in the model, semantic predictors have the strongest and most widespread influence on neuronal activity, indicating that meaning remains the dominant driver of neural responses in the hippocampus.

Next, we used the same linear modeling to predict neural activity from up to five words in the past and future. We found that neural responses to the current word could be predicted from word embeddings up to one word in the past and up to two words in the future (LLH differences were always significantly greater than 0, see **Methods**).

However, prediction accuracy showed a characteristic decay in decodability as a function of distance forwards and backwards in time (**Fig. 1g, right**). Thus, it appears that hippocampal neurons keep track of recently heard words and also show some evidence of encoding upcoming words, just as other brain regions have been shown to do (Goldstein et al., 2024; Goldstein et al., 2022; Goldstein, Wang, et al., 2023; Jamali et al., 2024; Tang et al., 2023; Zada et al., 2024). These findings suggest that hippocampal neuron responses integrate information across multiple words and could contribute to representing contextual semantics during natural speech. Note that these effects could also arise from overlapping neural responses across adjacent words or from correlations between nearby words in natural speech. Additionally, whether this future word encoding represents a true prediction is a debated issue; our results do not resolve that debate (Schönmann et al., 2025).

We next identified the broad semantic categories that exist in our dataset.

Word2Vec embedding distances recapitulate semantic distance, such that the embedding vectors for synonyms like ‘dad’ and ‘father’ have a smaller semantic distance (cosine distance) than unrelated words like ‘dad’ and ‘emergency’ (Joulin et al., 2016; Mikolov, Chen, et al., 2013; Mikolov, Sutskever, et al., 2013). We leveraged this fact to build semantic categories (**Fig. 1h**; Huth et al., 2016; Jamali et al., 2024). Silhouette criterion analysis (**Fig. 1i**; Rousseeuw, 1987) and k-means clustering of the word embeddings (see **Methods**) revealed 11 putative semantic categories in the story data, listed in increasing word frequency, with examples in brackets: (1) body parts [hand, lungs, scar], (2) places-[hospital, office, cabin], (3) emotional [cheer, disgust, feelings], (4) mental-[know, answer, imagine], (5) social [friends, queen, dad] (6) objects [book, flowers, pigeon], (7) visual [beautiful, green, sizzling], (8) numerical [another, eleven, moment], (9) actions [shimmied, writing, elected], (10) identity [she, y’all, their], and (11) function words [the, about, onto] (**Fig. 1j**). Note that the categorization here is based on a static Word2Vec embedding rather than a contextual embedding, such as that provided by BERT or GPT-2. In this case, clustering of single words based on contextual embeddings wouldn’t produce meaningful categories, because word meanings are determined by their local contexts. We show these categories in Extended Data Fig. 6a-b. We will cover contextual embeddings below (**Figs. 3-4**).

**Fig. 3.**
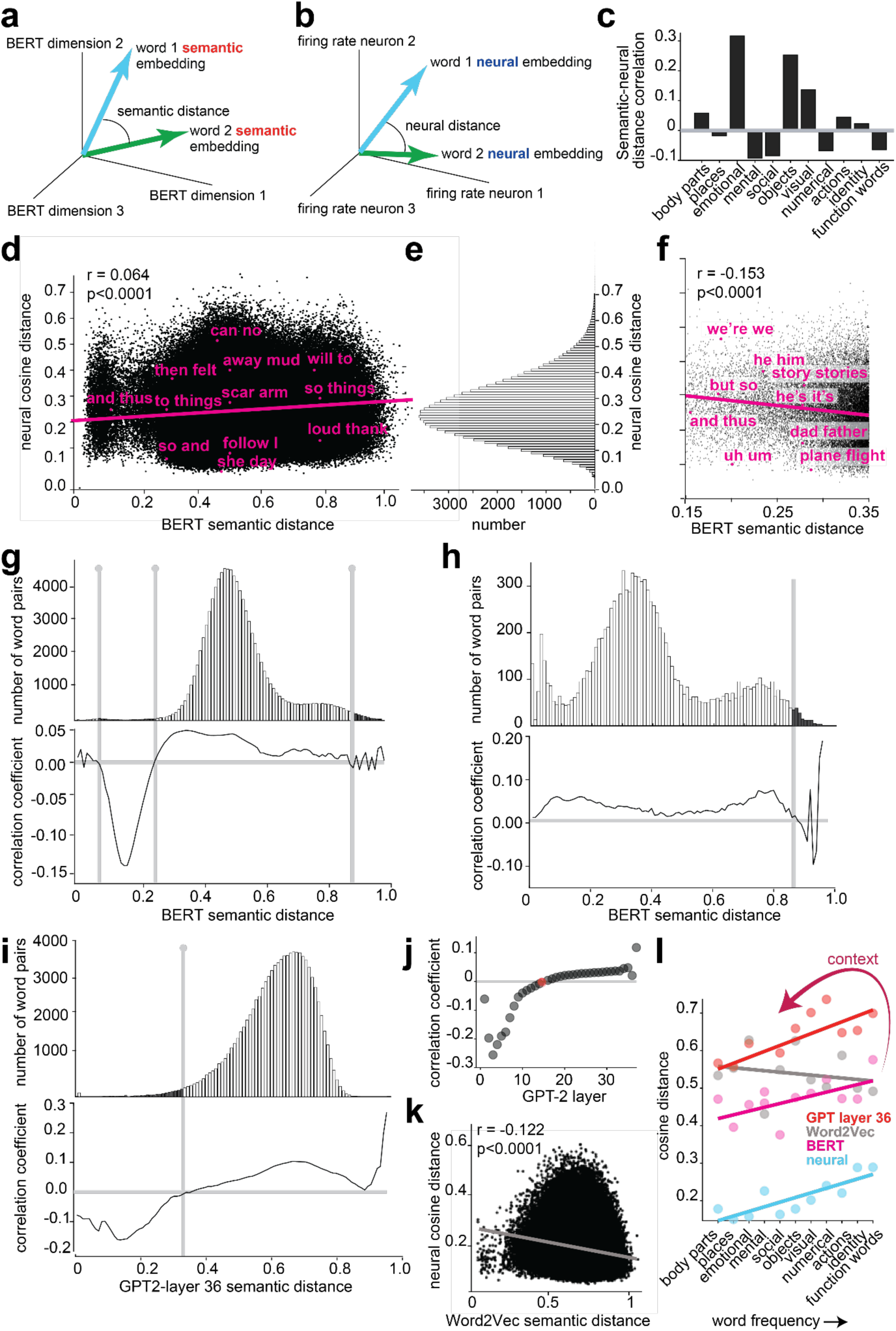
Neural distance recapitulates semantic distance. a, We define semantic distance as the angular distance in a high dimensional space between two semantic vectors (embedding values for each word). **b,** We define neural distance in the analogous manner, where each word’s neural embedding is the vector of firing rates from all neurons for that word. **c,** Semantic BERT embedding and neural embedding distance correlations among all word-pairs within each of the eleven semantic categories. While most are positive, some categories have negative correlations, likely due to contrastive coding. **d,** Scatter plot illustrating the correlation between neural and semantic embedding distances between all 26,978,185 word pairs. Pink line: regression line. Each dot represents one randomly selected word pair; magenta text indicates example pairs. **e,** Histogram of neural embedding distances; this plot is the marginal of **d** across the x-axis. **f,** Expanded view of the data from **d** in the neural range of 0.15-0.25; this range represents 2.16% of the data. In this range, the correlation between neural and semantic distances is negative. **g,** Histogram of BERT semantic embedding distances; this plot is the marginal of **d** across the y-axis. Below it is shown the semantic-neural correlation coefficient for the sub-range of BERT values centered on the x-axis point. While this line is generally positive in value, at especially short distances (low x-values), it turns negative. That negative correlation may reflect contrastive coding. **h,** Same data as in **g**, but limited to pairs in which the same word is compared in two locations. Because BERT is a contextual embedder, these will not have precisely the same value. **i,** Same data as in **g**, but using GPT-2 layer 37 rather than BERT. Same effects are observed, indicating that effects are not due to BERT. **j,** Correlation between semantic and neural distance across all layers of GPT-2. Early layers that are non-contextual embeddings have negative correlations with neural data, while higher, contextual layers are positive. Red indicates a layer with nonsignificant correlation. **k**, The same as in **d** but with Word2Vec semantic embeddings. These embeddings are static and non-contextual (similar to early layers in GPT-2), and negatively correlate with neural distance. **l,** Within-category neural distance and semantic distances from different language models. The arrow reflects how context changes semantic embeddings, improving their association to neural patterns of semantic coding. Word2vec r =-0.22, BERT r = 0.62*, GPT-2 r = 0.72*, Neural r = 0.83*; Pearson correlation coefficient, *P < 0.05, regression lines plotted.

**Fig. 4.**
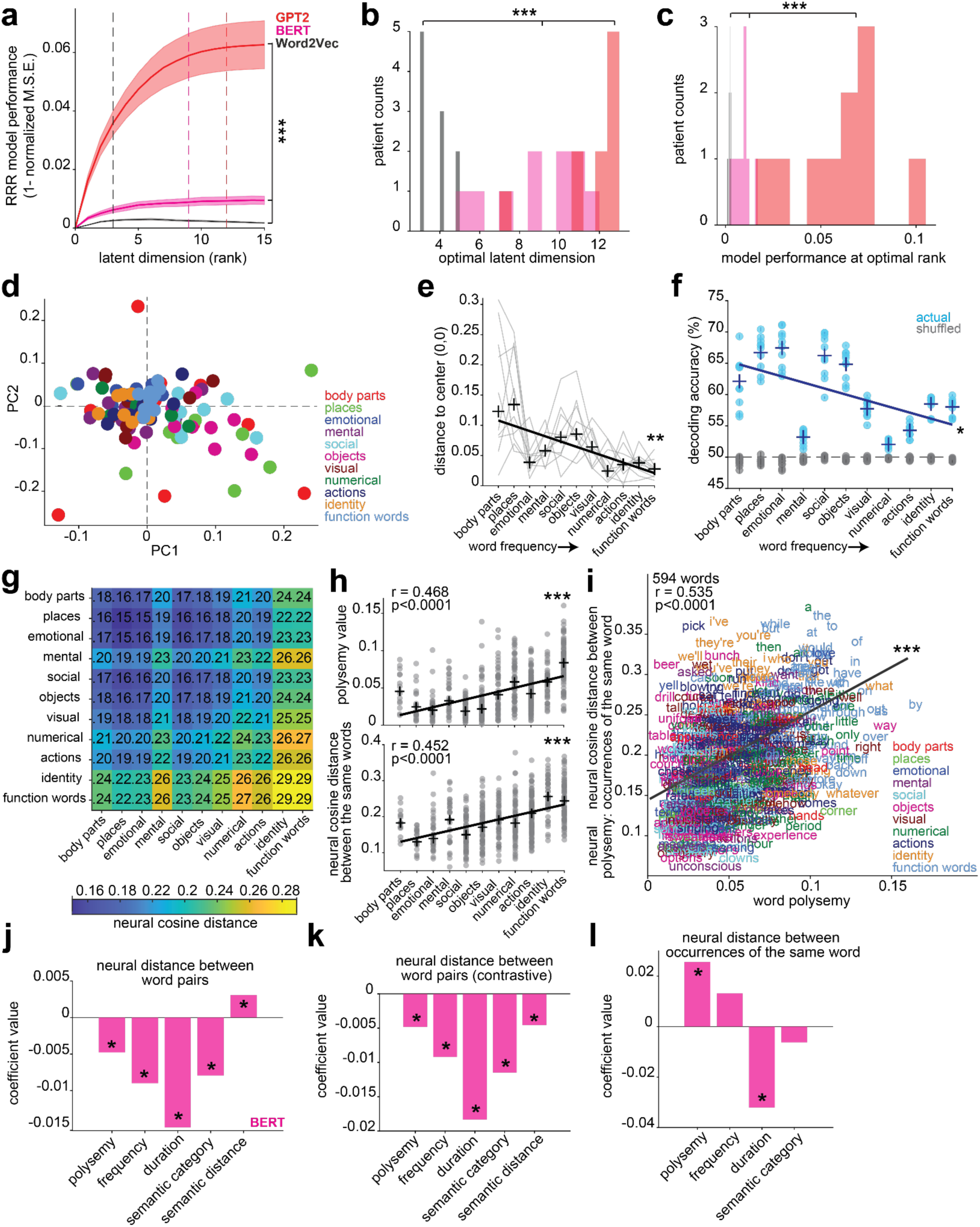
Population coding of words across frequency and context. a, Model performance across 15 latent dimensions (ranks) from reduced rank regression (RRR) analysis predicting semantic embeddings from neural firing rate responses to words. Models were fit separately for each patient, then averaged across. Lines represent mean performance ±SEM. Model performance was significantly different between models with Word2Vec, Bert, and GPT-2 embeddings. Asterisks in **a-c** represent P-values from a one-way ANOVA comparing metrics from GPT-2, BERT, and Word2Vec models. **b,** The optimal latent dimension for each patient and model. **c,** Model performance at the optimal rank for each patient and model. **d,** Centroids of words within semantic categories from each patient plotted in a 2D shared semantic subspace. **e,** Mahalanobis distance of each semantic category centroid to the origin (0,0), computed separately for each patient (gray lines); crosses represent median distance. Distance significantly decreases across semantic categories; r =-0.78, P = 0.004. **f**, For each patient, decoding accuracy for each semantic category and all other categories from neural population activity, for models using actual (blue) and shuffled (gray) labeled data. Chance is 50% (dashed line). Crosses represent patient averaged decoding accuracy. Plots display mean prediction accuracy on test observations (±SEM). P < 0.0001 for each category, Wilcoxon signed-rank test comparing actual and shuffled accuracy. Decoding accuracy significantly decreases across semantic categories; r =-0.54, P = 0.02. **g**, Similarity matrix of vectorial neural distance between all word pairs within and across each semantic category. The diagonal of this matrix is plotted in Fig. 3l. **h**, Top - polysemy values for each unique and repeated word within each semantic category. Bottom - neural distance between different occurrences of the same word, within each semantic category. P and r-values are from Pearson correlation. **i**, Each word is plotted according to its polysemy value and neural distance (median distance between all occurrences of that word). Words are colored based on their semantic category. P and r-values are from Pearson correlation. **j,** Weights from a ridge regression model that predicted neural distance between all word pairs from the difference in polysemy, word frequency, word duration, and semantic category for each word pair, and the BERT semantic distance between them. Asterisks represent significant predictors. **k,** The same as in **j** but restricted to word pairs with semantic distances within the contrastive coding range, 0.14 - 0.4. **l,** Weights from a ridge regression model that predicted, for every word, the median neural distance between all occurrences of that word from that word’s polysemy (computed with BERT embeddings), frequency, duration, and semantic category. Asterisks represent significant predictors. Lines in e, f, h, and i are regression lines.

### A distributed coding of semantics

One possible way to represent meaning in the brain is through neurons that selectively respond to single words, or to single categories of semantically related words, such as those associated with family, sports, or nature (Jamali et al., 2024). However, this categorical coding regime could be biologically inefficient, as it would require many neurons - enough needed for all the concepts learned across one’s life. We hypothesized instead that word coding would be highly distributed, with neurons exhibiting mixed selectivity-meaning one word should be encoded by many different neurons and a single neuron should encode many, semantically unrelated words (Rigotti et al., 2013; Fusi et al., 2016; Johnston et al., 2020 and 2024).

To quantify individual neuron encoding of words, we performed a Receiver Operating Characteristic (ROC) analysis (**Fig. 2a-b**), a standard approach that has previously been used to characterize neural responses during behavior (Britten et al., 1992; Li et al., 2017). For statistical reasons (**Methods**), we used only words that occurred at least four times in the data (n = 283 words), and compared firing rates evoked by each word to all others (see **Methods**). Every single one of these 283 words was encoded by at least 3 neurons (**Fig. 2c-e**). The words with the least encoding (i.e., most sparsely coded) were ‘was’ and ‘and’, encoded by exactly 3 neurons; the most dense word, ‘hit”, was encoded by 67 neurons (0.8 and 18.8% of our sample, respectively, **Fig. 2d-e**). On average, a single word is encoded by 37 neurons (10.4%, **Fig. 2d**), suggesting a distributed code for semantics. Likewise, most neurons encoded many words. Only 6 out of 356 neurons did not encode any words, and the median number of words encoded by a neuron was 6 words (**Fig. 2f**).

Are the words a neuron encodes semantically similar or different? Surprisingly, the same neuron does not reliably encode semantically similar words. For example, within the social category (category 5), we examined encoding of words related to the concept of family (mom, dad, father, brother, and son), and found minimal overlap between neurons that encode each family-related word (**Fig. 2g**). Indeed, of the 28 neurons encoding “father” and 19 encoding “dad,” only 7 encode both words (**Fig. 2g**). This low overlap reflects the contextual nature of hippocampal semantic coding during natural language comprehension - neurons encode the meaning of a word as it is used in a specific narrative context, rather than responding to membership in a broad semantic category. Notably, the distinct words that a cell encoded spanned semantic categories and phonology. For example, neuron 167 encodes the words: ‘lot’, ‘remember’, ‘even’, ‘they’, ‘those, and ‘make’; neuron 252 encodes the words: ‘she’, ‘last’, ‘out’, ‘school’, and ‘team’. Each of those neurons encode words that span 5 unique semantic categories. Overall, a given neuron’s response field spanned 4 of our 11 semantic categories (**Fig. 2i**). (And, recall that these proportions refer to all sampled neurons, not just a preselected subset of selective neurons).

Finally, for each neuron (limited to neurons that significantly encoded at least two words, n = 334), we quantified its ***semantic range*** by computing the cosine distance between embeddings of all pairs of words it encoded. We compared this distribution to a null one that represents the full semantic space of the word data (**see Methods).** On average across embedding models, 56% of neurons (186/334) encoded words that spanned the full semantic range of the data, as determined by both bootstrap-based significance (≥ 95% support) and 95% confidence-interval coverage criteria (see **Methods**). 91% of total neurons (304/334) encoded words that span at least 70% of the full semantic range. Together, these results demonstrate that hippocampal neurons exhibit broad, mixed semantic tuning.

We also wondered whether more specific concepts, like proper nouns, were encoded in this distributed manner. Only three proper nouns appeared frequently enough in our data to perform statistics: ‘Mark,’ ‘God,’ and ‘Michelle’. These three are also encoded across neurons in the population, with ‘God’ showing the highest density at 60 neurons (**Fig. 2h**). 89 neurons (25%) encoded these 3 proper nouns, with minimal neuronal overlap between them (14 neurons encoded all 3 names). Across semantic categories, fewer neurons encoded identity (pronouns) and function words, while more critical and less frequent categories, such as body parts and emotion words, were distributed - encoded by many neurons (**Fig. 2j**). We found that such densely coded words had significantly higher classification performance (area under the curve) compared to sparsely coded words, suggesting neurons exhibit more distinct firing rates patterns for these less frequent words that are critical for understanding (**Fig. 2K**).

### Hippocampal population activity reflects semantic relationships

In semantic embedders like Word2Vec, words are represented as points in a high dimensional space, and distance in this space corresponds to semantic distance (**Fig. 3a**). Thus, synonyms like “mom” and “mother” are located close to each other in embedding space, while unrelated terms like “mom” and “tarmac” are located far apart. Here, we follow the standard approach of using angular (cosine) distance between embedding vectors to quantify vectorial distance (Mikolov et al., 2013; Pennington et al., 2014; Radford et al., 2019). We performed an analogous analysis on the responses of a population of neurons to ask whether *neural embedding distances* are correlated with semantic embedding distances (**Fig. 3b**). Overall, we find that contextual embedding distance between randomly selected word pairs is correlated with their neural embedding distances (**Fig. 3c-e**). Consider, for example, the contextual embedder BERT (using unidirectional embeddings; Lyu et al., 2023; Madureira et al., 2024; Kahardipraja et al., 2021; Devlin et al., 2018; Pearson r = 0.0650, p<0.0001, **Fig. 3d**; Note that each data point represents the cosine distance (on the y-axis) of neural responses between two single words, with no averaging across repetitions.). Using GPT-2, we also found a significant positive correlation between neural distance and semantic distance (r = 0.0921, p<0.0001, **Extended Data Fig. 7a and Fig. 3j**).

Note that while the size of this correlation is low, this is inevitable given the extremely high variance of single neuron firing rates and the fact that each datapoint reflects a single occurrence of a given word pair. Indeed, we quantified the maximum correlation achievable between the actual semantic embeddings and their altered version that mirrors neural data (see **Methods**), and found that it was 0.122 and 0.119 for BERT and GPT-2 respectively. Therefore, the neural-BERT correlation of 0.06 captures 54% of the noise-limited ceiling, showing that over half of the model’s representational structure is recoverable despite the high variability shown by single neurons for single repeats of a word (**Extended Data Fig. 7d**). The neural-GPT-2 correlation of 0.09 is, relatively, even stronger, reaching 76% of the ceiling (**Extended Data Fig. 7b-d**). Moreover, neural-semantic correlations exceed the entire shuffle-based null distribution where neural responses were shuffled across words, indicating that the effect reflects true neural-semantic structure rather than an artifact of having many data points (gray histograms, **Extended Data Fig. 7d**).

This problem of high noise in neuron’s firing rates could cause confusion for words with the most similar embeddings. Such word pairs include “but and so”, “we and we’re”, and “he and him” (although the embeddings also include local context not given here, **Fig. 3f**). While the brain needs to recognize their similarity, it also needs to maintain their semantic separation as well. Thus, we hypothesized that, at the finest scale, neuronal embeddings would have an additional feature of contrastive coding (sometimes called repulsive coding; Bromley et al., 1994; Chanales et al., 2017; Prince et al., 2025; Radford et al., 2021; Tian et al., 2020), which would allow for separation between similar, but still distinct, words. We find precisely this: pairs of closely related words (regardless of semantic category) tend to show a *negative* correlation between semantic and neural distance (**Fig. 3f-g**). In other words, the relation between semantic and neural embedding is scale-dependent: negative for the finest scale and positive for all others (**Fig. 3g**). Specifically, the negative pattern is observed when considering the word pairs whose semantic distances span the range of r = 0.15 to r = 0.35 (these comprise 2.16% of the words in our sample, **Fig. 3f-g**). (Note that words with a range of r < 0.15 comprise less than one tenth of one percent of pairs, and data in this range are too noisy to meaningfully interpret). These highly similar pairs include many pairs of identical words; however, this negative correlation in semantically close pairs is driven entirely by word pairs consisting of different words (near-synonyms like “sort/kind” and “are/were”), rather than identical word pairs (such as “and/and” or “the/the”). Specifically, in the range of 0.15-0.35, we found r =-0.151 (p < 0.0001) for different word pairs (**Fig. 3f**) and r = +0.032 (p = 0.45) for identical pairs (**Fig. 3h**).

Notably, this coding regime was not unique to BERT embeddings. The same contrastive coding for semantically similar word pairs and pattern separation for semantically different words is also observed in GPT-2 layer 37 embeddings (notice the similarity between **Fig. 3g and 3i** and **Extended Data Fig. 7a-d**; Radford et al., 2019). Interestingly, we found a negative correlation between semantic and neural distance for early GPT-2 layers (1-13) and for Word2Vec embeddings (**Figure 3j-k**). We think this is due to the early layers of GPT-2 being less contextual than the late layers, and thus more similar to static, non-contextual embeddings from models like Word2Vec. Therefore, the neural coding of semantics is both vectorial and highly contextual. Note that all results in Figure 3 were replicated when performed with populations of neurons from each patient individually (**Extended Data Fig. 7e-g**). Additionally, removing function words and pronouns slightly increased neural–semantic distance correlations, suggesting that these words add context-driven variability while content words yield more consistent semantic responses (**Extended Data Fig. 5c**). Because this does not change our results qualitatively, we retain them in the analysis.

Interestingly, the distance between neural embeddings of words within a semantic category is a function of increasing word frequency (**Fig. 3l, blue**; this is the case for inter-category distances as well). Meaning, low-frequency body part words like “hand” and “finger” are closer in both neural space and the semantic space of contextual embeddings, while high-frequency identity words like “everyone” and “they” show greater distance, as seen in **Fig. 3l** (red, pink, and blue). In other words, frequent words are more centrally positioned in the high-dimensional embedding space, a pattern that is replicated in the hippocampal coding. Indeed, applying context to semantic embeddings causes a reduction in semantic distance for low-frequency words and increases it for high-frequency words, compared to static embeddings like Word2Vec (**Fig. 3l, gray**).

This shift, influenced by context, aligns with the pattern of category and frequency-dependent neural distance, perhaps explaining why Word2Vec embedding distances are negatively correlated with neural data, whereas contextual models show a positive correlation, more accurately reflecting the vectorial and contextual encoding of semantics in the hippocampus.

### Semantic coding is contextual

The meaningful correspondence observed between neural and semantic distances led us to test a central prediction: that a word’s semantic representation should be decodable from its neural population activity. We applied cross-validated reduced-rank regression to assess how well neural population activity predicted 30-dimensional semantic embeddings from Word2Vec, BERT, and GPT-2 (Izenman 1975; Semedo et al., 2019; Wu and Pillow, 2025). Model performance was quantified as the cross-validated variance explained (1 – NSE) across predictive subspace dimensionalities (ranks = 0 - 15). All models showed near-zero performance at rank 0 (predicting the mean), confirming an unbiased baseline. Across ranks, predictive performance increased monotonically before reaching a plateau. GPT-2 achieved the highest accuracy (patient-averaged 1–NSE = 0.06), followed by BERT (0.01) and Word2Vec (0.003), with a significant effect of model type **(Fig. 4a**; F(2, N) = 82.3, p = 9.3 × 10⁻¹⁶). Additionally, performance plateaued at different subspace dimensionalities for each language model: median rank of 12 for GPT-2, 9 for BERT, and 3 for Word2Vec (dashed lines, **Fig. 4a**). These optimal ranks reflect the dimensionality of the predictive neural subspace required to account for semantic structure in each embedding space. The higher optimal rank and superior performance for the contextual models (GPT-2 > BERT > Word2Vec) indicate that neural population activity shares a richer, more multi-dimensional linear structure with GPT-2 embeddings than with Word2Vec. This pattern implies that semantic information is distributed across multiple neural dimensions rather than confined to a single feature axis, consistent with a population-level, vector-like code. In contrast, the lower optimal rank for Word2Vec indicates that non-contextual embeddings capture a narrower, lower-dimensional component of the neural representation. The fact that GPT-2 and BERT outperform Word2Vec even at rank 1 suggests that the primary neural dimension is contextual, not categorical - if it reflected a single feature like ‘people vs. places,’ all models should perform similarly at rank 1 (or Word2Vec would have the highest performance) because the decoder is limited to expressing only that one contrast. Together, these results demonstrate that contextual language models provide a substantially better and higher-dimensional description of the semantic information expressed in single-neuron population activity. Findings were consistent across patients (**Fig. 4b-c**).

Given this evidence for a high-dimensional neural-semantic space, we next examined how semantic categories are arranged within this population geometry. For each patient separately, we performed Principal Component Analysis (PCA) of neurons’ firing rates for every word (7,346 words by patient’s neurons), reducing the dimensionality of their neurons’ responses to 2 components. To visualize neural organization of semantics across patients, we first projected each patient’s neural data into the first two principal components. These two components explained 44% of the total neural variance, on average. Next, we computed the centroids of each semantic category in PCs 1 and 2, and then aligned this data across patients into a common reference frame, yielding a shared low-dimensional subspace (see **Methods**). Within individual patient subspaces and within this shared one, we noticed the first axis (PC1) separated social, objects, and places, from identity, mental, and action words. Notably, there appeared to be an organization based on word frequency, where high-frequency words clustered near the center and low-frequency words were more spread out (**Fig. 4d**). Indeed, when we computed, for each patient, their semantic category centroid to subspace center distance, we found a significant decrease in distance with increasing word frequency (r =-0.78, p = 0.004; **Fig. 4e**), meaning high-frequency words were more overlapping and central in the subspace.

Notably, this representational geometry aligned with decoding performance, meaning semantic categories that were further from the center in the subspace were more easily decodable compared to geometrically clustered categories (r = +0.37, p = 0.0001, Pearson correlation). Each semantic category can be decoded from the others using population activity (see **Methods**), but high frequency words like numbers, actions, and function words had reduced decoding performance compared to low frequency words (**Fig. 4f**). Together, these findings suggest, as do others above, that neural coding principles differ markedly for high frequency and low frequency words.

Indeed, we also observe a trend in word frequency for neural distance. For word pairs within each semantic category, neural distance increases with word frequency (**Fig. 3l**, blue and **Fig. 4g**, diagonal), meaning neural firing rate population patterns between two common words are more distinct than patterns between two low frequency words. In fact, high-frequency words exhibited the greatest neural separation both among themselves and relative to words from other semantic categories (**Fig. 4g**). Although PCA projects high-frequency words close to the center (because their distinctions lie in low-variance axes not captured by PCs1-2, **Fig. 4d**), vectorial distance shows that they are broadly separated across the full high-dimensional neural space.

Why do common words cluster in neural state space, have reduced decoding, and increased vectorial neural distance (within-category separability) compared to other, less spoken words? We think this is due, in large part, to the fact that common words are more polysemous (**Fig. 4h, top;** Zipf, 1945), meaning the same word can have different meanings depending on the context in which it is spoken. To explore this idea further, we computed polysemy for each word by taking the cosine distance between BERT embedding vectors of repeated occurrences of the same word (n = 594 distinct and repeated words, see **Methods;** Cruse et al., 1986; Landauer, 2001; Ethayarajh, 2019). We then computed “neural polysemy”, or the neural distance between all occurrences of the same word, and for each word, took the Pearson correlation between its median neural distance and polysemy value. We find a strong significant correlation between neural polysemy and word polysemy (r = 0.54, **Fig. 4i**). This positive correlation remains even when function words and pronouns are removed (r = 0.49, **Extended Data Fig. 5d**).

We also computed polysemy for each word as the total number of synsets (i.e., distinct senses) associated with each word in WordNet (Fellbaum, 1998) and these polysemy values yielded the same results shown in Fig. 4h-i (Pearson r = 0.514, p < 0.001). Polysemy is likely driving the larger neural distance between words occurring *within* high-frequency categories compared to low-frequency ones (**Fig. 4g and 3l**). The words within these categories have more nuanced and flexible meanings, resulting in greater semantic (in contextual models) and neural distance within categories (**Fig. 4g and 3l**).

Polysemy also explains why high frequency categories are harder to decode from all others - because these common words activate broad, overlapping neural patterns rather than category-specific ones. For example, ‘the’ may produce similar neural patterns to the noun that it modifies, such as ‘the school’, making ‘the’ and ‘school’ harder to neurally distinguish, despite being members of two different semantic categories (function words and places, respectively). Indeed, when words are clustered using neural embeddings rather than semantic embeddings, the resulting groups do not align with clear semantic categories, consistent with the idea that word meaning in the brain is highly context-dependent (**Extended Data Fig. 8**). Given that neural distance positively correlates with semantic distance in contextual models (**Fig. 3**), it follows that neural activity patterns also strongly correlate with polysemy (**Fig. 4h, bottom and 4i**). In other words, the meaning of a given high-polysemy word is more contextually driven than the meaning of a low-polysemy word, and hippocampal encoding tracks with that contextually driven meaning.

Because multiple linguistic features covaried with neural distance and polysemy, we next asked which of them primarily drove the observed changes in neural activity patterns. For each word pair, we used semantic distance, differences in duration, frequency, and polysemy, and a binary indicator of whether the words belonged to the same semantic category (0 = same, 1 = different) as predictors in a cross-validated ridge regression model to predict neural distance (see **Methods**). We found that all predictors were statistically significant, but semantic distance (from contextual models such as BERT or GPT-2, see **Extended Data Fig. 9**) was the only one with a positive coefficient, meaning that greater semantic dissimilarity was associated with greater neural separation (**Fig. 4j**; semantic distance results were reversed in Word2Vec, see **Extended Data Fig. 9**). Limiting the model to the contrastive-coding range (semantic distance 0.14 - 0.4) produced a sign reversal for semantic distance, whereas all other predictors remained negative (**Fig. 4k**). This behavior identifies semantic distance as the sole predictor whose relationship to neural geometry systematically tracks the emergence of contrastive coding. Additionally, we used the same model architecture to predict neural polysemy from word polysemy, word frequency, word duration, and semantic category. We found that only word polysemy and duration were significant predictors of neural polysemy, with polysemy associated with greater context-driven neural variability and longer duration associated with more stable neural responses (**Fig. 4l**). Repeating the polysemy and related analyses using only neurons that remained semantically selective after controlling for other linguistic features produced similar results, consistent with a distributed population code for semantics, where semantic relationships are carried by patterns of activity across many neurons rather than by a small group of highly selective units (**Extended Data Fig. 10**).

## DISCUSSION

Here, we show that the hippocampus exhibits a distributed and contextual coding scheme for semantics. That is, individual words are encoded in the simultaneous activity of multiple neurons whose selectivities span multiple unrelated semantic categories (**Fig. 2**). While a single neuron can distinguish individual words, the semantic *relationship* between words (even multiple occurrences of the same word) is reflected in neural population activity (**Figs. 3-4**). This pattern is reflected in the association between neural activity and embeddings derived from high layers of contextual embedders (BERT and GPT-2), which incorporate meaning from adjacent words and sentences. These results, then, support the idea that neuronal coding of word meanings is similar to that observed in word models and LLMs, in which single vector entries do not correspond to specific concepts, and may not, on their own, have natural interpretations. Instead, meaning emerges from the collective pattern of activity across the population.

In many prior studies, semantics has been operationalized in terms of discrete lexical categories (e.g., actions, animals, or people; Jamali et al, 2024; Huth and Gallant, 2016). However, our results suggest that hippocampal neurons encode aspects of contextual semantics - how a word’s meaning is modulated by its linguistic context - rather than fixed lexical semantic categories during natural language comprehension.

Notably, the distributed semantic coding and mixed selectivity in single hippocampal neurons we observed appears to contrast the semantic coding regime in the left posterior middle frontal gyrus (pmFG), where neurons respond selectively to specific semantic domains (Jamali et al., 2024). Most pmFG neurons (30%) were selective for words related to actions, and most neurons that exhibited such semantic selectivity (84%) were selective for one semantic domain and no others (Jamali et al., 2024). Both that study and ours examined semantic coding as participants listened to natural sentences. Why would single neurons in two different brain regions, the pmFG and the hippocampus, encode semantics so differently? We theorize pmFG favors selective coding of literal word meanings because of its proximity and anatomical connectivity to speech production regions, whereas the hippocampus exhibits a flexible encoding of relational and context-dependent semantics, given its role in memory. Extending single-neuron studies to other brain areas will reveal how broadly - and in what form - semantics is encoded across the human brain.

Surprisingly, at short semantic distances (∼0.14-0.4), we find a less expected effect, in the form of an anti-correlation between neural responses and linguistic embeddings for the most similar 2-4% of words. We propose that this reversal reflects a contrastive code that allows the brain to distinguish semantically very similar words (**Fig. 3f and Extended Data Fig. 7e,g**). That is, for similar words (like “*house*” and “*home*”), the coding is more different than would be expected given the pattern observed with randomly sampled words. Contrastive coding appears to be driven by the semantic embedding distance between two words and not differences in their other linguistic features like changes in word duration, word frequency, and word polysemy, or whether the two words belong in the same semantic category (**Fig. 4j-k**). Contrastive coding is observed in all of our patients individually (**Extended Data Fig. 7e,g**), and is highly statistically significant. This is not a pattern that is observed in word model or large language model embeddings, so it may reflect constraints imposed by the brain. One such constraint is neural noise - highly similar embeddings may be more likely to be confused with each other in the brain than *in silico*. Yet, highly similar embeddings may be crucial to distinguish. For example, “*brother*” and “*sister*” have very similar embeddings, as do “*hamburger*” and “*veggieburger*” and “*eighteen*” and “*nineteen*”. However, these words do not have precisely the same meaning, and there are occasions when it is important to make the kinds of fine distinctions that they require (e.g., when buying birthday gifts, when feeding a vegetarian friend, and when paying for a parking spot). Likewise, a contrastive code may help with fine distinctions in context; for example, a word like “*generous*” or “*fun*” may have both sincere and ironic meanings, or even intermediate (shaded) meanings. Notably, confusing these word meanings may not be a problem in word models (like word2vec and LLMs) which may have very different noise properties than the human brain. Thus, the brain may use a distance-based approach at most scales, but would add additional contrastive elements to the most confusable words, and rely on a complementary approach for assessing similarity, at the finest scales. Such a complementary approach may include a combination of high shattering dimensionality and high generalization, which allows simultaneous separation and combination (Bernardi et al., 2020). That contrastive code would appear in our data as the pattern we observe; however, follow-up studies, likely with more carefully controlled datasets, will be needed to fully test this speculative hypothesis.

Another striking result is that, even in our models incorporating many linguistic features, semantic predictors exert the strongest and most pervasive influence on neuronal activity - even though the variance explained by semantic embeddings, or their predictability in decoding analyses, is modest in absolute terms (26% variance explained in encoding models, 1-6% variance explained in population decoding models). A key consideration for interpretation is that our naturalistic design fundamentally differs from paradigms that rely on repeated presentations of discrete stimuli, as commonly used in most experimental paradigms that study language processing in single neurons (Dijksterhuis et al., 2024; Xu et al., 2025; Jamali et al., 2024; Khanna et al., 2024). Indeed, our stimuli are not presented many times in an identical form, but instead are embedded in continuous, context-rich narrative speech and occur only once. In natural language, the meaning of a word is strongly context-dependent, and averaging over repetitions would inappropriately collapse distinct contextual states. Consequently, modest explained variance is expected. Our study adds to a small but growing literature using semantic embeddings and encoding models to investigate the neural representation of semantics. Prior work relating semantic representations to neural activity during naturalistic language has reported similarly modest effect sizes across different neural signals, including BOLD responses (Huth & Gallant 2016) and intracranial field potentials (Goldstein et al. 2024; Zada et al., 2024; Goldstein et al. 2022). Related work using human single hippocampal neurons responding to visually presented stimuli has also reported comparable relationships between neural activity and semantic similarity, although the specific metrics and experimental paradigms differ from those used here (Karkowski et al. 2025). Importantly, when accounting for the intrinsic sparsity and Poisson-like variability of single-unit activity, our results (**Fig. 3**) represent a substantial fraction of the noise-limited ceiling for semantic relationships between words: for example, the neural-semantic distance correlation of 0.06 for BERT reaches 54% of its estimated ceiling, and GPT-2 reaches 76%. These findings demonstrate that, even under the demanding constraints of single-presentation naturalistic stimuli, neural population activity captures a surprisingly large proportion of the recoverable semantic structure, supporting the view that modest absolute variance can still reflect robust, theoretically meaningful relationships.

Why are semantic embedding distances anti-correlated with neural embedding distances for Word2Vec? Our data do not provide a clear answer to this question, but, in aggregate, they suggest the outlines of an answer. Overall, we find that contextual embedding distances and neural embeddings both align with word frequency and polysemy levels (which themselves are highly correlated, **Fig. 3l**). Thus, the meaning of a high-polysemy word is more influenced by context than that of a low-polysemy word - this makes sense because high polysemy words, almost by definition, have meanings that are highly influenced by context. We suspect that it is this context that is the primary driver of the correlation between neural and semantic embeddings. The non-contextual (static) embedders lack all of this context and give words a single context-free embedding. Absent the context, the underlying overall negative correlation is uncovered. This underlying negative correlation does not have a ready explanation, but may reflect a special property of the hippocampus, which is associated with contrastive coding (Bromley et al., 1994; Chanales et al., 2017; Prince et al., 2025; Radford et al., 2021; Tian et al., 2020). Note that, contrary to our findings, another study shows a positive correlation between single neuron and semantic non-contextual embeddings in pmFG, highlighting the distinct coding repertoire used by the hippocampus (Jamali et al., 2024).

If hippocampal semantic coding is distributed, then it shares key representational features with semantics in LLMs. This does not imply that the brain functions like an LLM, and our data highlight clear differences between the two systems - for example, anticorrelations between neural and embedding distances for non-contextual embeddings and for very close words. Rather, the parallel lies in their use of high-dimensional, distributed population structure: individual units (neurons or embedding dimensions) are not interpretable on their own, yet distances in the population space reliably reflect semantic relationships. This shared geometry does not suggest a shared algorithm; it indicates that both biological and artificial systems can converge on similar representational organization. LLMs therefore provide a useful comparative framework for interpreting neural population structure, not because they mirror the brain’s mechanisms, but because they offer a high-dimensional representational model against which neural data can be contextualized.

## METHODS

### Human intracranial neurophysiology

All experiments were performed in the Epilepsy Monitoring Unit (EMU) at Baylor St. Luke’s Hospital using standardized approaches (Xiao et al., 2024A and B).

Experimental data were recorded from 10 adult patients (6 males and 4 females) undergoing intracranial monitoring for epilepsy. The hippocampus was not a seizure focus area of any patients included in the study. Single neuron data were recorded from stereotactic (sEEG) probes, specifically AdTech Medical probes in a Behnke-Fried configuration. Each patient had an average of 3 probes terminating in the left and right hippocampus. Electrode locations are verified by co-registered pre-operative MRI and post-operative CT scans. Each probe includes 8 microwires, each with 8 contacts, specifically designed for recording single-neuron activity.

Single neuron data were recorded using a 512-channel *Blackrock Microsystems Neuroport* system sampled at 30 kHz. To identify single neuron action potentials, the raw traces were spike sorted using the *Wave_clus* sorting algorithm (Chaure et al., 2018)and then manually evaluated. Noise was removed and each signal was classified as multi or single unit using several criteria: consistent spike waveforms, waveform shape (slope, amplitude, trough-to-peak), and exponentially decaying ISI histogram with no ISI shorter than the refractory period (1 ms). The analyses here used all single and multiunit activity (see **Extended Data Fig. 1**).

### Electrode visualization

Electrodes were localized using the software pipeline intracranial Electrode Visualization (iELVis; Groppe et al., 2017) and plotted across patients on an average brain using Reproducible Analysis & Visualization of iEEG (RAVE; Magnotti et al., 2020). For each patient, DICOM images of the preoperative T1 anatomical MRI and the postoperative Stealth CT scans were acquired and converted to NIfTI format (Li et al., 2016). The CT was aligned to MRI space using FSL (Jenkinson et al., 2002; Jenkinson & Smith, 2001). The resulting coregistered CT was loaded into BioImage Suite (version 3.5β1; Joshi et al., 2011) and the electrode contacts were manually localized. Electrodes coordinates were converted to patient native space using iELVis MATLAB functions (Yang et al., 2012) and plotted on the Freesurfer (version 7.4.1) reconstructed brain surface (Dale et al., 1999). Microelectrode coordinates are taken from the first (deepest) macro contact on the Ad-Tech Behnke Fried depth electrodes. RAVE (Magnotti et al., 2020) was used to transform each patient’s brain and electrode coordinates into MNI152 average space. The average coordinates were plotted together on a glass brain with the hippocampus segmentation and colored by patient.

### Natural language stimuli

Patients with epilepsy and healthy language function listened to six episodes (5- to 13-minutes in duration) taken from *The Moth Radio Hour,* and totaling 47:25 minutes of listening time (7,346 words). The six stories were, “Life Flight”, “The Tiniest Bouquet”, “The One Club”, “Wild Women and Dancing Queens”, “My Father’s Hands” and “Juggling and Jesus”. In each story, a single speaker tells an autobiographical narrative in front of a live audience. The six selected stories were chosen to be both interesting and linguistically rich. Stories were played continuously through the built-in audio speakers of the patient’s hospital television. The audio signal was synchronized to the neural recording system via analog input going directly from the computer playing the audio into the Neural Signal Processor at 30 kHz.

### Jabberwocky stimuli

After story listening, a subset of four patients listened to 17 minutes of Jabberwocky stories (1,890 words) to assess neural specificity for semantics (**Fig. 1d and Extended Data Fig. 2b-c**). Jabberwocky stimuli are words that are altered from the real words in the stories used in the experiment to become words that have no meaning, for example, “toom” or “skook”. An example sentence of jabberwocky stimuli is: Before I smow it, the foodbace toom is at my frose woor, and a coakle of them have bapes of bour under their alk. We used the software, *Wuggy*, to transform story transcripts into jabberwocky (Keuleers & Brysbaert, 2010). This program matched the subsyllabic segment length of each word and replaced words with letters that match phonetic probabilities of orthographic English. Jabberwocky preserves syntactic structure by retaining function words and morphemes, even though it lacks coherent semantics.

### Audio Transcription

After experiments, the audio.wav file was automatically transcribed using Python and Assembly AI, a state-of-the-art AI model to transcribe and understand speech. The transcribed words and corresponding timestamp output from Assembly AI was converted to a TextGrid and then loaded into Praat, a software for speech analysis. The original.wav file was also loaded into Praat and we manually checked the spectrograms and timestamps, correcting each word to ensure the word onset and offset times are accurate. The TextGrid output of corrected words and timestamps from Praat was converted to a.xls and loaded into Matlab and Python for further analysis.

### Semantic embedding extraction from language models

***1) Word2Vec.*** We used the pre-trained *fastText Word2Vec* model in MATLAB to extract word embeddings for all words in our dataset (Joulin et al., 2016; Mikolov et al., 2013). This pre-trained model provides 300-dimensional word embedding vectors, trained on millions of words text, to capture semantic relationships between words. Any surname words, such as “Harwood” or proper nouns like “Applebee’s” that did not have word embeddings were discarded from the analysis (these were rare in our sample).
***2) GPT-2 Large***. In an excel file of our transcribed words, each row corresponded to an individual token, including punctuation and sentence terminators such as “.”, “?”, and “!”. We reconstructed sentences by iterating through the file and appending words until encountering a recognized sentence delimiter. This process yielded multi-word segments that more accurately captured natural linguistic boundaries. Any remaining words following the last delimiter were grouped into a final sentence. Notably, punctuation tokens were preserved to maintain contextual fidelity but were tracked carefully to avoid introducing sub-word alignment errors during tokenization.

To extract GPT-2 embeddings, we employed the “gpt2-large” model (Radford et al., 2019) from the Hugging Face Transformers library (Wolf et al., 2020). This version of GPT-2 features a total of 37 layers (1 initial embedding layer plus 36 transformer layers), each producing 1280-dimensional hidden states, with a maximum context length of 1024 tokens.

Although GPT-2 supports a 1024-token context window, all embeddings used in this study were computed within individual sentences only - that is, the model’s context was reset whenever a sentence-ending delimiter (e.g., “.”, “!”, “?”) was encountered. We extracted hidden states from all 37 layers using a progressive context approach designed to incorporate GPT-2’s autoregressive nature. Specifically, for each sentence, we began with an empty context and added words sequentially. After adding each new word, we tokenized the updated text string via GPT-2’s Byte Pair Encoding (BPE) tokenizer, ran the token sequence through the model in no-grad (inference-only) mode, and retrieved the hidden states. We then identified the newly appended word’s sub-word tokens and averaged their corresponding vectors to obtain a single word-level embedding. Because GPT-2 often splits words into multiple sub-word fragments (otherwise known as “tokens,” e.g., “computa” + “tion”), averaging those fragments preserves consistency with the transcript’s original word boundaries. The GPT-2 outputs resulted in 1280-dimensional embeddings for each layer, as seen in **Fig. 3j**. All analyses with GPT-2 used word embeddings from the last layer 37, unless otherwise specified, as in **Fig. 3j**.

***3) BERT.*** We conducted a parallel extraction procedure with the “bert-base-cased” model (Devlin et al., 2018), also accessed via the Hugging Face Transformers library (Wolf et al., 2020). In contrast to GPT-2, whose progressive context reflects its decoder-only design, BERT uses a masked-language-modeling objective and an encoder-only architecture that processes all input tokens in parallel, granting access to both past and future context. To align BERT’s outputs more closely with the unidirectional processing of GPT-2, we applied an expanding-context procedure: beginning with an empty string, we appended one new word at a time, concatenated the running text (preserving punctuation for contextual fidelity), and re-tokenized the updated segment with BERT’s tokenizer. This process continued until a sentence-ending delimiter (e.g., “.”, “!”, or “?”) was encountered, at which point the context was reset, ensuring that embeddings were computed strictly within sentence boundaries. Although BERT inherently considers bidirectional information, we extracted only the hidden states corresponding to the just-added word, thereby restricting interpretation to left-contextual information only. Sub-word vectors belonging to that word were averaged to yield one word-level embedding per layer. Hidden states from all 13 layers (768 dimensions each) were retained, and analyses reported in this paper used the final hidden layer 13. All code was implemented in Python using PyTorch (Paszke et al., 2019) as the underlying deep-learning framework, and the resulting embeddings were saved to Excel spreadsheets for subsequent analyses.

### Clustering of semantic embeddings

To identify the natural semantic categories present in our word data (**Fig. 1h-j**), all unique words were clustered into groups based on related meanings using a word embedding approach (Joulin et al., 2016; Mikolov, et al., 2013). For each word, we used its 300-dimensional Word2Vec embedding. To compute and visualize semantic clusters, we used a t-distributed Stochastic Neighbor Embedding (t-SNE) algorithm on word embedding values to reduce the dimensionality of each unique word based on their cosine distance to all other words, thus reflecting their semantic similarity. Words with similar meanings now have similar 2D coordinates. We then applied the K-means clustering algorithm to these 2D word representations and visualized clustered words on a 2D word map (k = 11 clusters, determined from the silhouette score; Rousseeuw, 1987). We then manually inspected and assigned a distinct semantic label to each cluster and adjusted categories for accuracy. For example, words bordering the edges of categories might get mis-grouped and were manually corrected. This method is comparable to those used in other studies of semantics (Huth et al., 2016; Jamali et al., 2024). We repeated this same method to create semantic clusters from neural data (**Extended Data Fig. 8**) and contextual embeddings (**Extended Data Fig. 6**).

### Firing rate responses to words

We computed the duration for each word in the story and for each non-word in Jabberwocky stimuli. The firing rate of a neuron for each word was the number of spikes occurring during the word with an 80 ms delay after word onset to account for the approximate delay of information to hippocampus. This value was divided by word duration and then multiplied by 1000 to compute spikes per second.

### Regressing spike counts on word embeddings, phonemes, and linguistic features

Word embeddings, such as those derived from Word2Vec or similar models, are typically high-dimensional and can exhibit substantial collinearity among dimensions - that is, many of the embedding features are correlated with one another. We first used Principal Component Analysis (`pca`, Matlab 2023b) on the full word embeddings to obtain uncorrelated features that still capture the dominant structure in the embedding space with reduced dimensionality. For each word, we used the first 100 principal components (PC) from Word2Vec or BERT embeddings because they explained at least 60% of the variance of the original embedding vectors.

We modeled the spike count responses of individual neurons using a Poisson Generalized Linear Model (GLM) with a log-link function and ridge (L2) regularization (we confirmed that spike counts for each word follow a Poisson distribution; **Fig. 1f-g**). The model aimed to predict the number of spikes a neuron fired in response to each word (summed spikes across word duration), using the top 100 principal components (PCs) of the word embedding vector, the duration of the word, and the interaction between each PC and the word’s duration as predictors (z-scored prior to model fitting). This resulted in a 201-dimensional feature vector for each word: 100 PC values, 1 duration value, and 100 PC×duration interaction terms.

The Poisson GLM assumes that the spike count *y_i_* for word *i* is drawn from a Poisson distribution with mean λᵢ where:

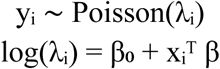

Where 𝜆_i,n_, the expected spike count for word (*i)* and neuron n (*n*), is modeled by the linear regression coefficients (𝛽), one for each of the 201 predictors (xᵢᵀ), plus the y-intercept (𝛽_0_). For each neuron, we used a 5-iteration nested training loop, where in each iteration, the data was split into training and held-out test sets (80/20 split). Within the training set, 5-fold cross-validation was used to select the optimal regularization strength (alpha) from a log-spaced range based on optimizing the cross-validated log-likelihood. The selected model was then evaluated on the held-out test set using several performance metrics, including log-likelihood, and adjusted R². To assess statistical significance relative to a null, we generated 100 shuffled predictor matrices per dataset, in which embeddings were shuffled while keeping word duration preserved. Each shuffle circularly shifted the embedding matrix along the observation axis by a randomly selected offset of at least ±100 samples (to avoid short-range autocorrelation artifacts). To determine if the actual model showed significant improvement from the shuffled, we performed a permutation test comparing the actual model LLH to the shuffled model LLH distribution. We then took the difference between the actual and shuffled model (mean) LLH (LLH diff; ΔLLH). A neuron was fit by the model if its ΔLLH was both positive and significant and if McFadden’s pseudo R² value was positive (P < 0.05, permutation test, **Fig. 1g, left**). Shuffled model R² values were significantly less than those from actual models (p < 0.05, Wilcoxon signed-rank test; **Extended Data Fig. 4c**).

Predictor significance was assessed using p-values from a Poisson generalized linear model fit to the same predictors, while coefficient magnitudes were taken from the L2-regularized Poisson regression model used for prediction. As expected, word duration was always a significant predictor, followed by the first eight embedding dimensions and their interactions with duration, with the number of significant predictors gradually decreasing across higher dimensions (**Extended Data Fig. 4a-b, g-h**).

We used the same Poisson linear ridge regression model to predict neural activity from past and future word embeddings (**Fig. 1g, right**). Here, a separate model was fit for each word index, by which we shifted the words relative to the spike counts, such that the spike count of a neuron for word X corresponded to an embedding of a different word and word duration that was in the future or past, based on the index. To determine whether neural activity was predicted by each word index, the distribution of actual model performance (ΔLLH) from every neuron was compared to that of a null model (chance distribution, described above). For each word index, these distributions were compared using the one-sided Wilcoxon signed-rank test (p < 0.05).

Model performance was significantly greater than chance for up to seven words in the past and three words in the future.

### Phonetic regression

We built a fixed-length feature space over the 43 IPA phonemes observed in our corpus. Here, IPA refers to the International Phonetic Alphabet, a standardized set of symbols that denote speech sounds independent of spelling. The podcast transcript was processed word by word: tokens were lower-cased and converted from graphemes to phonemes using the Python phonemizer package (version 3.0.1) with the eSpeak backend as the grapheme-to-phoneme (G2P) model (https://pypi.org/project/phonemizer/3.0.1/). For each word, we took the canonical IPA transcription returned by phonemizer. Phonemes were then manually inspected and corrected by linguists (JB and BJ) in Praat.

Each word was then represented as a 43-dimensional count vector whose entries reflect the number of occurrences of each IPA phoneme in that word (unigram counts; no duration or stress information). This approach is similar to Khanna et al., in that words are represented by the presence of their constituent phonemes; however, we did not group phonemes into broader classes, in order to preserve the full identity of the 43 phones observed in our data and maintain the embedding’s informational richness and dimensionality (Khanna et al., 2024). To better differentiate words that contain the same phoneme set but in different orders (e.g., dog vs god), we double-counted the first phoneme in every word while counting all subsequent phonemes once. The resulting word-by-phoneme matrix (N words × 43 features) was used in subsequent analyses relating phonetic composition to neural responses. We used this word-by-phoneme matrix, word duration, and their interactions in the same Poisson regression described above for predicting spike counts of the current word. The data folds used for training and testing were preserved across semantic and phonetic models and both models had the same number of words. The results in **Extended Data Fig. 4e-f** were replicated even when using the same number of predictors (the first 43 principal components of semantic embeddings compared to 43 phonemes).

We used the same Poisson ridge regression model to predict an individual neuron’s spike count for each word from 61 linguistic features: (1) full phonetic embeddings (43 IPA phones; count coded, cf. Khanna et al., 2024), (2) 10 PCs of semantic embeddings, (3) word order in sentence, (4) word order in clause (Stanford CoreNLP; Manning et al., 2014), (5) opening node number (Stanford CoreNLP; Manning et al., 2014), (6) closing node number (Stanford CoreNLP; Manning et al., 2014), (7) word frequency (SUBTLEX-US), (8) pitch, (9) a scalar representing degree of polysemy (Cruse et al., 1986; Landauer, 2001; Ethayarajh, 2019) and (10) word duration (**Extended Data Fig. 4g-h**).

### Single neuron encoding of words

Semantic encoding of each neuron was quantified using a ROC (receiver operating characteristic) analysis, a standard approach that has previously been used to characterize neural responses during behavior (Britten et al., 1992; Li et al., 2017). Here, the firing rate of each neuron is the classifier that is evaluated on its ability to distinguish instances of a word from all other words. For statistical reasons, since we need to compare two distributions, we performed the analysis on every neuron for every word that occurred at least 4 times (283 words, on average each word occurred 20 times, the most was 407).

Upon application of a binary threshold to firing rates and comparison with a binary event vector denoting word label (1 for the current word, 0 for other words), word detection based on neural activity can be measured using the true positive rate (TPR) and the false positive rate (FPR). Plotting the TPR against the FPR over a range of binary thresholds, spanning the minimum and maximum values of the neural signal, yields an ROC curve that describes how well the neural signal detects that word at each threshold. With the `perfcurve` function in Matlab 2023b, we used the area under the ROC curve (AUC) as a metric for how strongly neurons are modulated by each word (**Fig. 2a-c**). The number of word occurrences in the ‘Current word’ class were balanced with the number of words in “Other” class by subsampling words from the “Other” class over 100 iterations of the analysis. For each neuron, AUC was computed, then averaged and compared to a null distribution across iterations. For each neuron, the observed AUC was compared to a null distribution of 1,000 AUC values generated from constructing ROC curves over randomly permuted neural signals (that is, firing rate permuted across words, thus destroying the relationship between neural responses and word labels). The p-value is calculated as the proportion of permuted AUC values that are greater than or equal to the observed AUC and a neuron was considered significantly responsive if P < 0.05. To better visualize encoding magnitude, we standardized each AUC value by subtracting chance (0.5) from every value, and took the absolute value of that difference. This results in values between 0 and 0.5 (**Fig. 2c**).

### Semantic range

Each neuron encodes a given set of words determined from ROC analysis (see *Single neuron encoding of words*). For each neuron that encoded at least two words, we defined its semantic range as the distribution of cosine distances between all pairs of words it encoded. We then compared this empirical range to that neuron’s null semantic range. For the null distribution, we randomly sampled the same number of encoded words from the full set of words used in ROC analysis from the dataset (n = 283 words), computed all pairwise cosine distances, and repeated this 1,000 times for that neuron. Thus, each neuron’s null distributions matched their empirical distribution count, representing samples from the full (testable) semantic space of the data. We also repeated this analysis using words and their embeddings from the full dataset (n= 7,346 words) and found comparable results.

We quantified the extent to which each neuron’s empirical semantic range encompassed the null range. Using nonparametric bootstrap resampling (n = 1,000), we generated distributions of the minimum and maximum for both the empirical and null ranges, and computed 95% confidence intervals (CI) from the bootstrap samples. A neuron’s semantic range was considered to cover the null range if the lower CI bound of its minimum was less than or equal to the upper CI bound of the null’s minimum, and the upper CI bound of its maximum was greater than or equal to the lower CI bound of the null’s maximum. Bootstrap support for coverage was defined as the proportion of bootstrap iterations meeting this criterion. Coverage was deemed significant if support was ≥ 1 − α (α = 0.05). A neuron was classified as spanning the full semantic range if it met both the CI-based coverage criterion and the bootstrap-significant coverage criterion. CI-based coverage identifies neurons whose observed range is statistically compatible with fully spanning the null range at the 95% confidence level, whereas bootstrap-significant coverage identifies neurons whose range covered the null range in ≥ 95% of bootstrap resamples. This analysis was performed separately using Word2Vec, GPT-2 (layers 36 and 25), and BERT (last layer) semantic embeddings.

### Embedding distance

The semantic embedding vector for each word was its full vector of embedding values extracted from either Word2Vec (300 in length), BERT (768 in length), or GPT-2 (1280 in length) language models. For each word, it’s neural embedding vector was every neuron’s firing rate response for that word (356 in length). Neurons were combined across all 10 patients, making a pseudo population totaling 356 neurons. We used the function `pdist`, Matlab 2023b, to compute cosine distance between semantic embedding vectors of all 26,978,185 word pairs, known as semantic distance. We repeated the procedure, computing cosine distance between neural embedding vectors for all word pairs, termed neural cosine distance. “All word pairs” included pairs of the same word except in Word2Vec analysis, since cosine distance between two instances of the same word will always be 0 (since embedding values are the same). This procedure was followed for word pairs within and between each semantic category to create the semantic distance similarity matrix in **Figure 1h**, for the neural and semantic distances within each category in **Figure 3l**, and for the neural distance similarity matrix in **Figure 4g**. For **Figure 3c**, we computed the semantic distance from BERT embeddings for all word pairs within each semantic category, then computed neural distance for all word pairs within each semantic category and correlated (Pearson) these two distance vectors to compute the values shown on the y-axis. Similarly, we correlated semantic distance and neural distance from all word pairs (unless otherwise specified in the caption) to compute the correlation coefficients in **Figure 3g-j**. In **Figure 4h-i**, neural cosine distance was always computed between word pairs of multiple instances (minimum 2) of the same word. We took the median of these distances to extract one distance value per word.

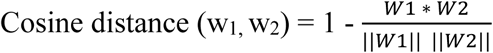

Cosine distance between neural or semantic embeddings of word 1 and word 2 is the 1 minus the dot product of embedding vectors for each word, divided by the magnitudes (norms) of each word vector.

### Estimating a neural noise ceiling for semantic embeddings

To estimate a noise ceiling constrained by the variability observed in the neural data, we generated noisy versions of the GPT-2 (or BERT or Word2Vec) embedding space by injecting Gaussian noise whose magnitude was scaled according to the firing-rate statistics of the recorded neurons. Let 𝐺 ∈ 𝑅^W×D^ denote the clean GPT/linguistic embedding matrix (rows = words, columns = embedding dimensions), and 𝑅 ∈ 𝑅^W×D^ the neural firing-rate matrix (rows = words, columns = neurons).

Because we empirically confirmed that hippocampal firing rates across words follow Poisson-like statistics, with each neuron’s variance closely matching its mean, we used a Poisson noise model in which the variance-to-mean ratio was fixed at 𝜙 = 1 for all neurons. This provides a conservative, interpretable baseline for quantifying how neural-level variability would distort the representational geometry of the language-model embeddings. Then, we injected additive Gaussian noise into the embedding space using a subset of embedding dimensions equal in number to the recorded neurons (D = 356).

For each word 𝑤 and embedding dimension 𝑑:

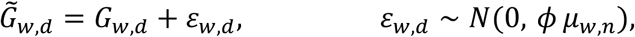

where 𝜇*_w,n_* is the mean firing rate of neuron 𝑛 (taken from 𝑅) corresponding to embedding dimension 𝑑, and 𝜙 = 1. Thus, the additive noise variance scaled linearly with the neurons’ mean firing rates, emulating Poisson variability across words. This model assumes that noise enters independently of the embedding magnitude, perturbing all points in embedding space by a constant variance level proportional to the corresponding neuron’s mean firing rate.

For each noise model, pairwise cosine-distance matrices were computed (using the function, `pdist`, in Matlab 2023b) for the noisy language embeddings (𝐷*_noisyGPT_*), the original language embeddings (𝐷*_GPT_*), and the neural data (𝐷*_neural_*). The noise-limited ceiling was defined as the Pearson correlation between 𝐷_noissyGPT_and 𝐷_GPT_:

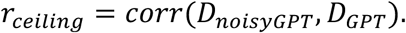

The observed brain–model correspondence was computed as:

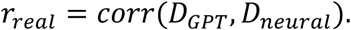

The ratio 𝑟_real:_/𝑟*_celling_*_<_quantifies how much of the model’s representational structure is recoverable in the neural data, relative to the theoretical maximum allowed by intrinsic neural variability.

### Reduced Rank Regression

To quantify how hippocampal population activity predicts high-dimensional semantic embeddings of heard words, we applied Reduced Rank Regression (RRR) using the Matlab code and following the framework of Semedo et al. 2019 (see also Wu and Pillow, 2025). For each patient, neurons served as predictors and individual words as observations (**Fig. 4a-c**).

Neural data preprocessing: For each patient, we computed per word firing rates normalized by word duration (see *Firing rate responses to words).* The resulting matrix contained one observation per word and one column per neuron (X). Word-level confounding variables - the log of word frequency and position - were regressed out of the neural activity matrix, and the residuals were used in analysis.

The first 30 Principal Components of semantic embeddings from Word2Vec, BERT layer 13, or GPT-2 layer 37 were used in the regression, forming a word by semantic embedding matrix (Y). RRR estimates a mapping from neural activity X to semantic embeddings Y that is constrained to lie in a low-dimensional shared subspace. We used the Semedo et al., 2019,

ReducedRankRegress function, which implements the closed-form solution for:

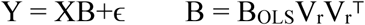

where 𝑉_r_ are the top 𝑟 eigenvectors of 𝑌^T^𝑋𝑋^T^𝑌. For each patient, we computed:

𝐵___: the ranked predictive components

𝑉: the eigenvectors describing predictive dimensions

These define the semantic subspace captured by the neural population. To determine the number of predictive dimensions, we evaluated models with 0–15 RRR dimensions using 10-fold cross-validation, following Semedo et al. Model performance (averages across folds) was quantified as the cross-validated variance explained (1 – NSE) across predictive subspace dimensionalities (ranks = 0 - 15). The optimal dimensionality (rank) can be defined as the number of dimensions needed for the prediction accuracy to plateau, selected using the Semedo et al., 2019 function, `ModelSelec`t. These RRR results quantify how many dimensions of semantic variation can be linearly predicted from hippocampal population activity, as well as the geometry of the neural to semantic mapping.

### Principal Component Analysis

We computed components (scores) from `pca`, MATLAB 2023b, on the word by neuron firing rates matrix for each patient (neuron count ranges from 11-54 per patient). To visualize and compare semantic representations across patients, we first constructed a common two-dimensional subspace (**Fig. 4d**). For each patient, word-by-neuron firing rate matrices were reduced with PCA and the first two principal components (PC1–2) were retained. Because the scale and covariance structure of these axes can vary across patients, we whitened each patient’s PC1–2 scores using their within-patient covariance matrix. This whitening procedure ensures that Euclidean distances in the transformed space are equivalent to Mahalanobis distances in the original coordinates.

Semantic category centroids were then computed as the mean of the whitened word representations belonging to each category. To align patients into a common reference frame, we applied an orthogonal Procrustes analysis to the configuration of category centroids, restricting the transformation to a pure rotation (no reflection, scaling, or translation). This yielded a shared coordinate system in which all patients’ category centroids could be visualized and compared.

Within this common space, we quantified the distance of each category centroid to the origin (0,0). Because whitening centers each patient’s space at their mean response, the origin corresponds to the population-average baseline. Distances were computed as the Euclidean norm of the whitened coordinates, which is equivalent to the Mahalanobis distance to the patient’s mean response in the original PC1–2 space. These values provide a standardized measure of how strongly each semantic category deviates from the center of the shared semantic subspace.

Results in Figure 4d-e were replicated using Euclidean distance between centroid and center (0,0) within each patient’s individual 2D subspace.

### Support Vector Machine Decoder

We used a SVM decoder (Bishop, 2006) with a nonlinear radial-based function kernel to determine whether firing rates during words carry information about semantics by distinguishing each category from all others (**Fig. 4f**). We also repeated this analysis with a linear kernel, and we found equivalent results to Figure 4f except performance was much lower, only 2-5% above chance level (results not shown). For each patient, each semantic category, we used the firing rate responses from every neuron (n = 11-54 across patients) for each word in the category (class 1) and all rates for all other words (class 2) to predict the class for each word from these neural data. The number of words in each class were always balanced – across comparisons and across semantic categories through random subsampling of words from the larger class. Random selections of class observations were repeated for 100 iterations, giving us the average classification accuracy over 1,000 test splits of the data for each session. To train and test the model, we used a tenfold cross-validation. In brief, the data were split into ten subsets, and in each iteration the training consisted of a different 90% subset of the data; the testing was done with the remaining 10% of the data. We used the default hyperparameters as defined in `fitcsvm`, MATLAB 2023b, with hyperparameter *C* = 1.3 and z-scored normalization of firing rates. Decoder performance was calculated as the percentage of correctly classified test trials.

We compared model performance for predicting train and test data to check for overfitting. In each iteration, we trained a separate decoder with randomly shuffled class labels. The performance of the shuffled decoder was used as a null hypothesis for the statistical test of decoder performance.

*Decoding jabberwocky stimuli.* We used the same methods described above on a linear SVM model (default hyperparameters) to decode whether a word was a real word from podcast stories or a non-word from the jabberwocky stimuli from neural population activity (**Extended Data Fig. 2c**). We excluded the function words from jabberwocky stimuli in this analysis since those words are real words but used in a nonsensical context.

### Polysemy

To compute polysemy for each word, we computed cosine distance between each BERT embedding vector of all occurrences of a given word (at least 2 occurrences, 594 words total).

We took the median of this distribution to obtain one polysemy value per word (**Fig. 4h-i**). The distribution of polysemy values for all words in each semantic category is plotted in Figure 4h – top. We repeated the polysemy analysis using neural embeddings instead of BERT embeddings to compute the neural cosine distance values in Figure 4h – bottom and Figure 4i. We also computed polysemy for each word as the total number of synsets (i.e., distinct senses) associated with each word in WordNet (Fellbaum, 1998). These polysemy values yielded the same results shown in **Figure 4h-i**.

### Modeling neural distance and polysemy from linguistic features

We used the same cross validated, linear Ridge Regression architecture described in *Regressing spike counts on word embeddings, phonemes, and linguistic features* to predict neural distance or neural polysemy from different linguistic features. Models fit in **Figure 4j-k** used word-pairs as observations and semantic distance, differences in duration, frequency, and polysemy, and a binary indicator of whether the words belonged to the same semantic category (0 = same, 1 = different) as predictors in a cross-validated ridge regression model to predict neural distance. In **Figure 4l**, the model used each word that was repeated at least twice as observations, and the word’s polysemy, frequency, duration, and semantic category (as defined by Word2Vec: categories ranged 1-11; treated as a categorical), were predictors for neural polysemy of that word. Ridge regularized coefficients/weights from these models are plotted in **Figure 4j-l**.

## Extended Data Figures

**Extended Data Fig. 1.**
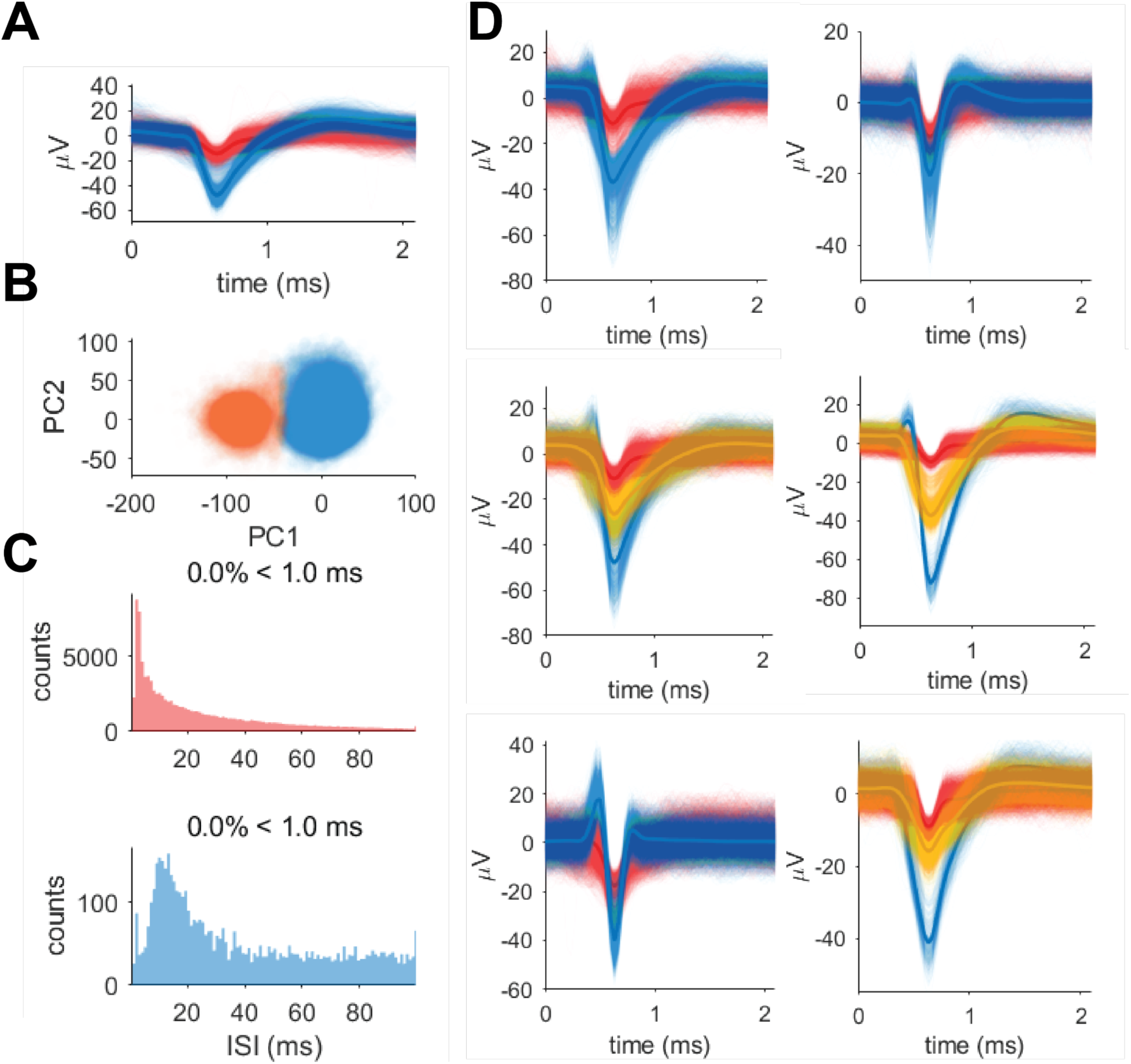
Single and multi-unit detection and isolation. A, A random sample of 5,000 individual waveforms from a single hippocampal neuron (blue) and multi unit cluster (red) from patient 8 recorded on one electrode channel. Bolded lines represent average waveforms. Noisy signals were already removed using *Wave_clus* sorting algorithms and manual inspection. **B**, All waveforms from each cluster in **A** are well isolated in Principal Component space. **C**, Interspike interval (ISI) histograms from the neurons in **A**. Single and multi-unit clusters always have refractory period violations below 5% (as shown in figure titles). **D**, Example hippocampal single (yellow and blue) and multiunit (red) waveforms on individual electrode channels from four patients that were identified and sorted following the criteria in **B-C** using *Wave_clus* software and manual inspection.

**Extended Data Fig. 2.**
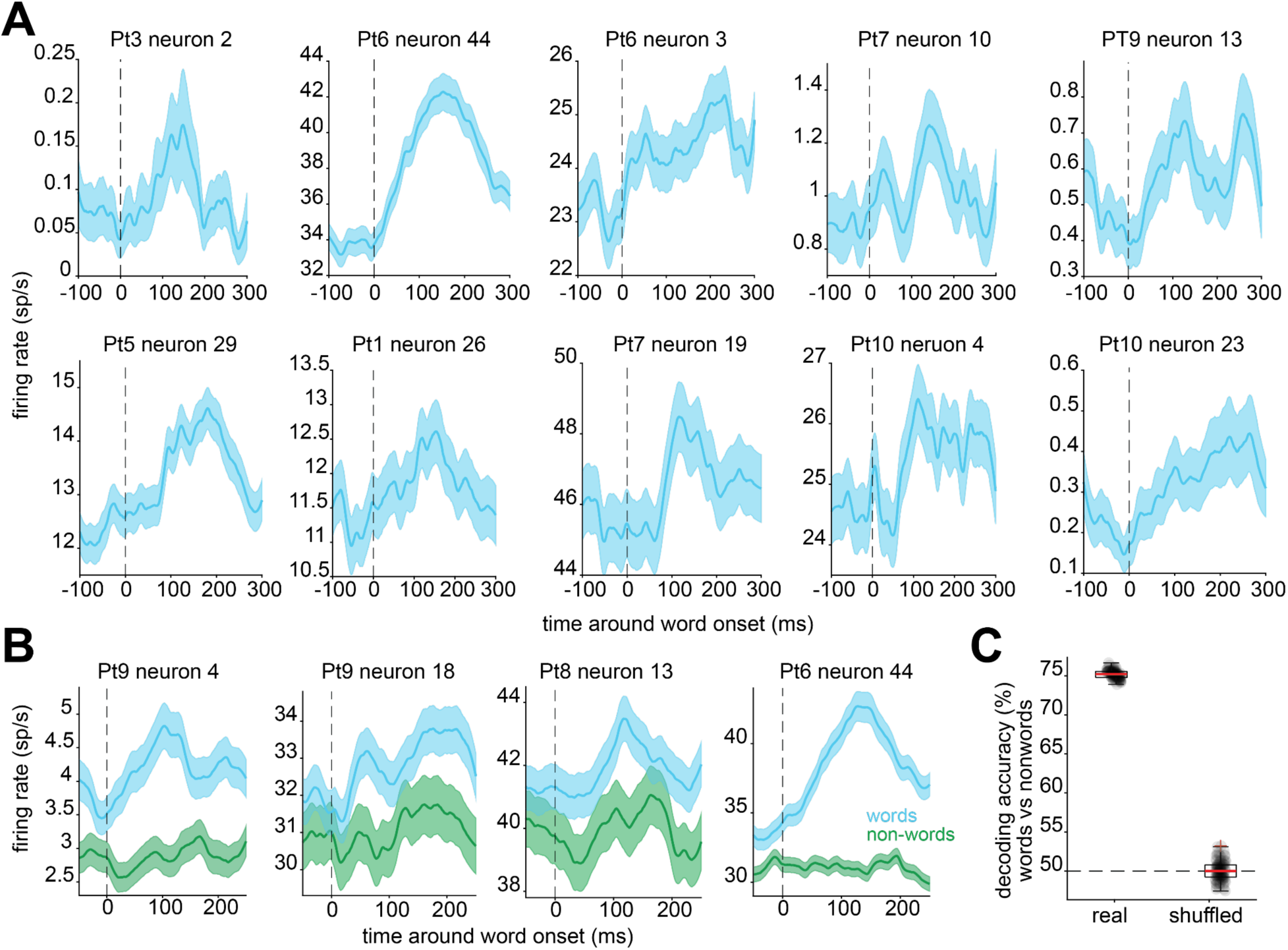
Hippocampal neuron responses to words and non-words. A, Peri-stimulus time histograms of firing rate responses to words from example hippocampal neurons in seven different patients during language comprehension. **B**, Peri-stimulus time histograms of firing rate responses of neurons exhibiting significant changes in firing rate to words and non-words (Wilcoxon rank-sum test, P < 0.05). Example hippocampal neurons are shown from 3 different patients. **C,** Linear decoding accuracy for words and nonwords (jabberwocky stimuli) from neural population activity comprised of all neurons across four patients, for models using real and shuffled data.

**Extended Data Fig. 3.**
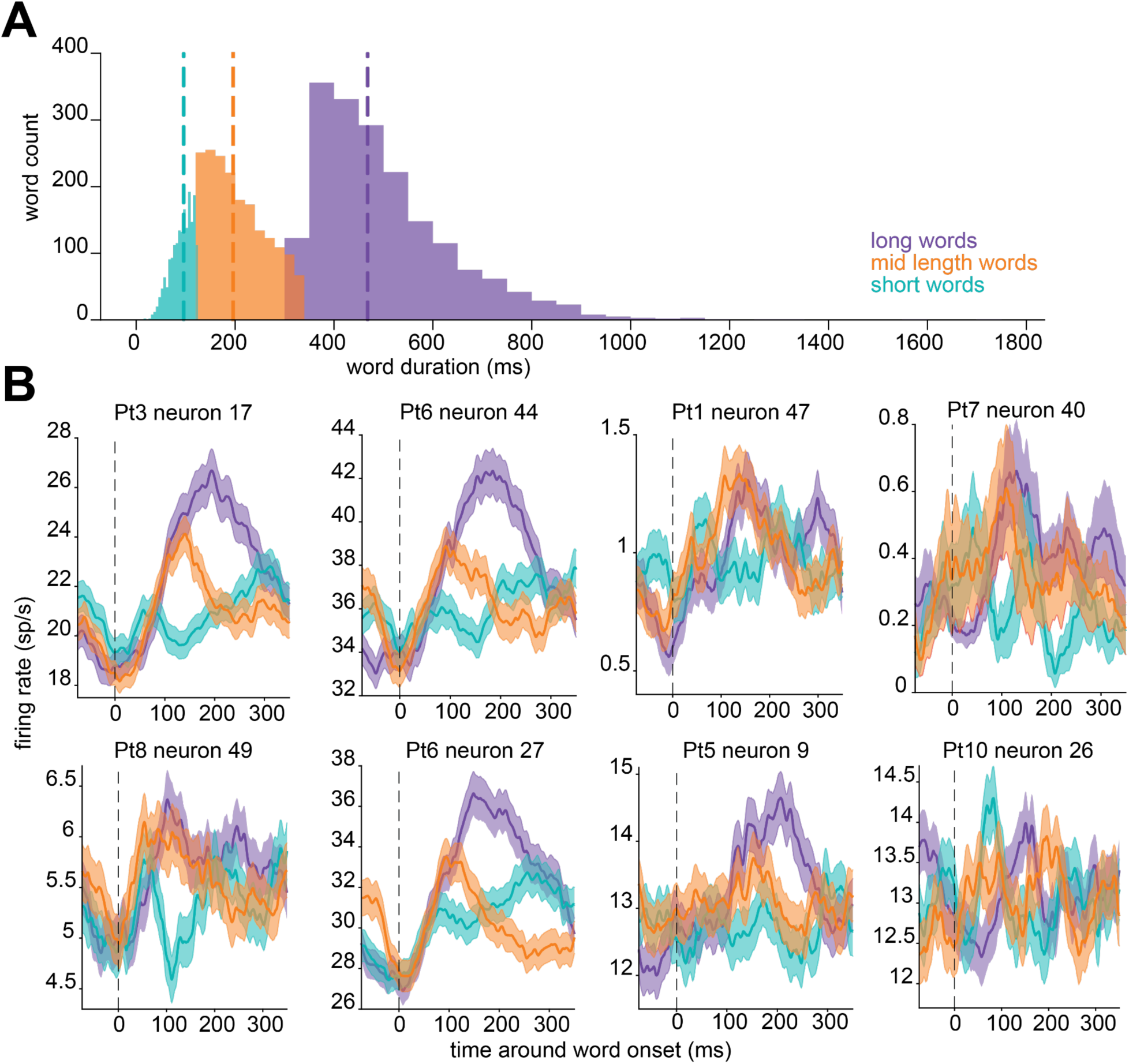
Single neuron responses to words of different spoken durations. A, Histograms of the word durations for short (1,852 words), middle length (1,852 words randomly sampled to match other groups), and long words (1,842 words). Dashed lines represent the median word duration for each group. **B,** Peri-stimulus time histograms of firing rate responses to words of different durations from example hippocampal neurons in seven different patients during language comprehension. Dashed lines represent word onset.

**Extended Data Fig. 4.**
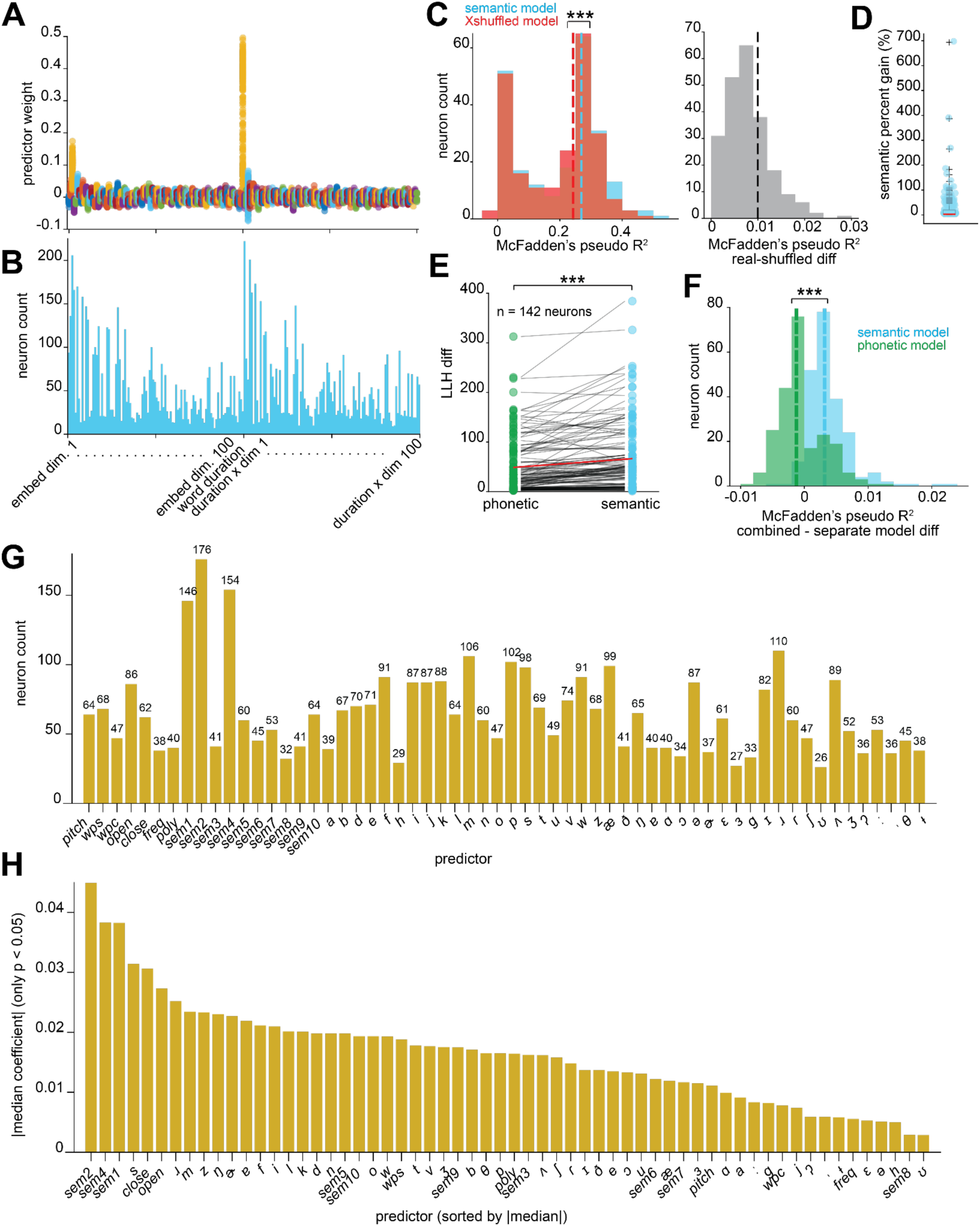
Semantic, phonetic, and combined linguistic feature regression models. A, Semantic model (Word2Vec) coefficients of significant predictors of neurons fit by the model. Word duration and embeddings 1-8 had the largest absolute valued weights. **B**, The number of neurons for which each predictor (as seen in **A**) was significant. PC embeddings 1-8, word duration, and their interactions were significant predictors in most neurons fit by the semantic model. **C,** Left - McFadden’s pseudo R-squared for neurons fit by the actual and embedding shuffled (X-shuffled) semantic model. Values from the real semantic model are significantly higher than shuffled; Wilcoxon signed-rank test, P < 0.001. Right-The difference in neurons’ Pseudo R-squared values between the semantic and X-shuffled model distributions (left). **D,** For each neuron, the relative increase in pseudo-R² when semantic embeddings are added to a word duration-only model. Specifically, it is R² of the combined full model minus R² of the duration model, divided by R² of the duration model and multiplied by 100. **E**, Log likelihood difference between real and shuffled models for the same neurons fit by the phonetic and semantic embedding models (see Methods); Wilcoxon signed-rank test, P < 0.001. **F,** For each neuron, the difference in pseudo R-squared from a combined model with phonetic and semantic embeddings, and a separate model with either phonetic (green) or semantic (blue) embeddings. Semantic embeddings explain significantly more variance than phonetic embeddings. Wilcoxon signed-rank test, P < 0.001. **G,** The number of neurons for which each predictor was significant in the model is plotted on the y-axis, and the total number of neurons is also displayed on top of the bar plot. This model used semantic embeddings from Word2Vec, and models using BERT or GPT-2 embeddings had comparable results. **H,** The absolute value of the coefficient, or weight, of each predictor in the model is shown on the y-axis. Bars reflect the median predictor weight taken across neurons fit by the model. Predictors are sorted by decreasing weights. Poly = polysemy; Wps = word position in sentence; wpc = word position in clause; freq = word frequency; open and close represent opening and closing nodes; sem1-10 are semantic embeddings from Word2Vec.

**Extended Data Fig. 5.**
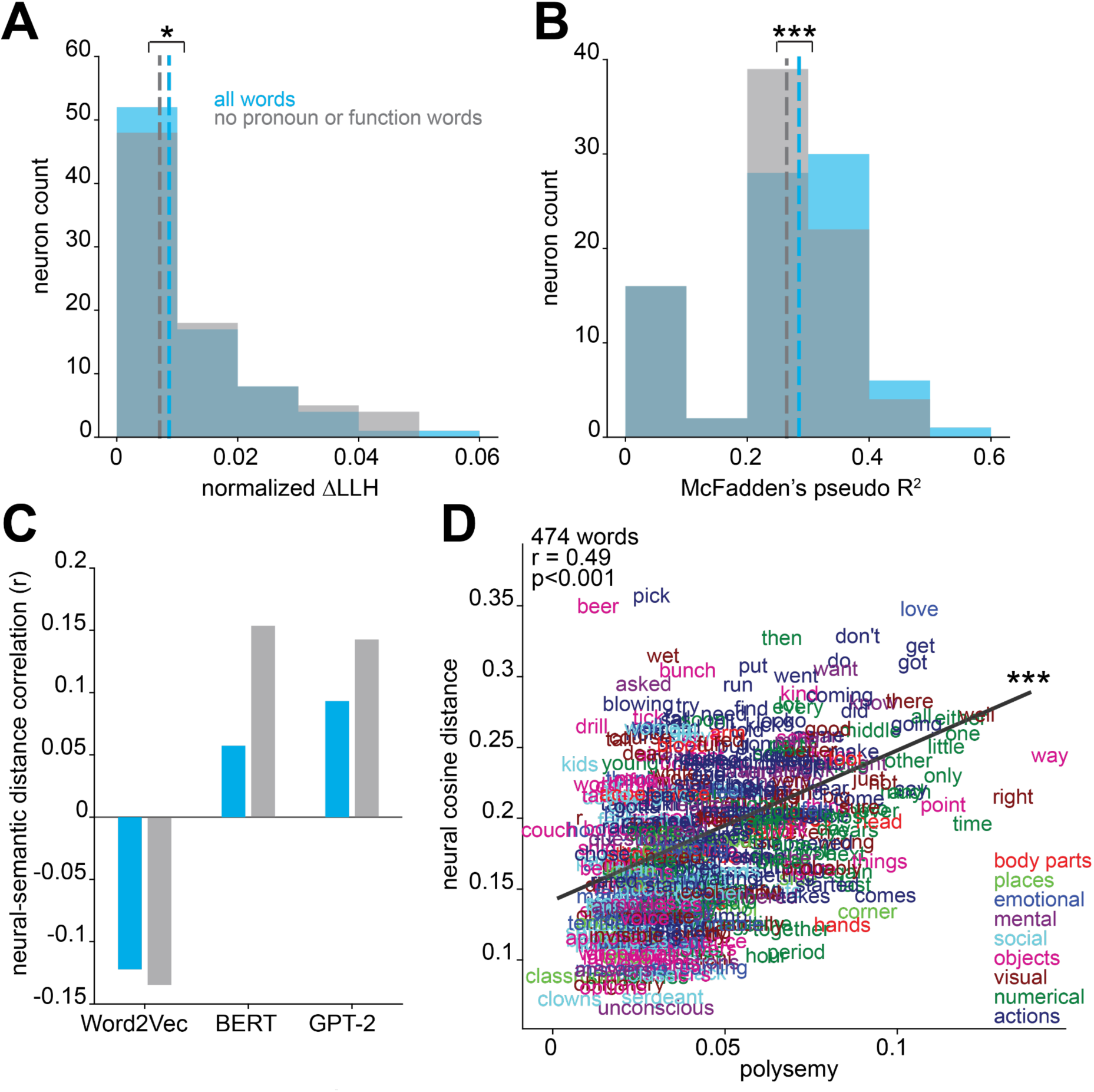
Effects of function words and pronouns on semantic coding. A, The change in log-likelihood between the null and actual model for models fit with all words (blue) compared to the same measure in models fit without function words or pronouns (gray, FWE model). Delta LLH values were normalized by dividing each neuron’s delta LLH value by the number of words used in the model. P< 0.05, Wilcoxon signed-rank test. **B,** The Pseudo R-squared values for the same neurons shown in A from models using all words or function words and pronouns removed. P< 0.001, Wilcoxon signed-rank test. **C,** The Pearson correlation coefficient between neural and semantic distance for either all word pairs (blue) or word pairs excluding function words and pronouns (gray). All correlations are significant, P < 0.001. **D,** Each word is plotted according to its polysemy value and neural distance (median distance between all occurrences of that word). Words are colored based on their semantic category. P and r-values are from Pearson correlation.

**Extended Data Fig. 6.**
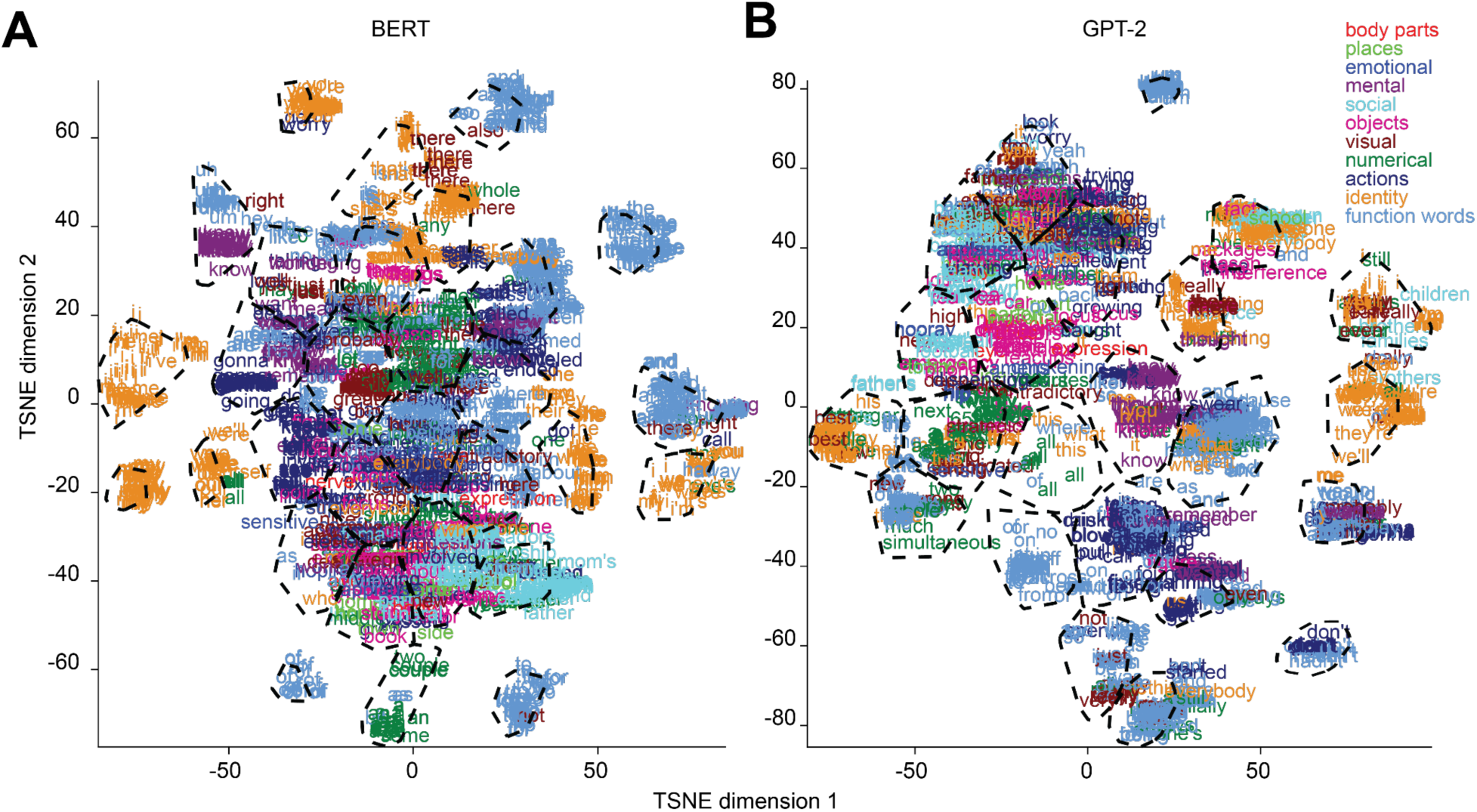
Semantic categories from contextual embeddings. A, words plotted in their dimensionality reduced BERT embedding space and colored according to their semantic category defined in Word2Vec. Words are grouped in 30 distinct categories defined from silhouette score and k-means clustering of BERT embeddings. Black dashed lines outline the identified categories. **B**, The same as in A but with GPT-2 layer 37 embeddings. 23 categories are identified. 1,808 words out of 7,346 are plotted here for visualization.

**Extended Data Fig. 7.**
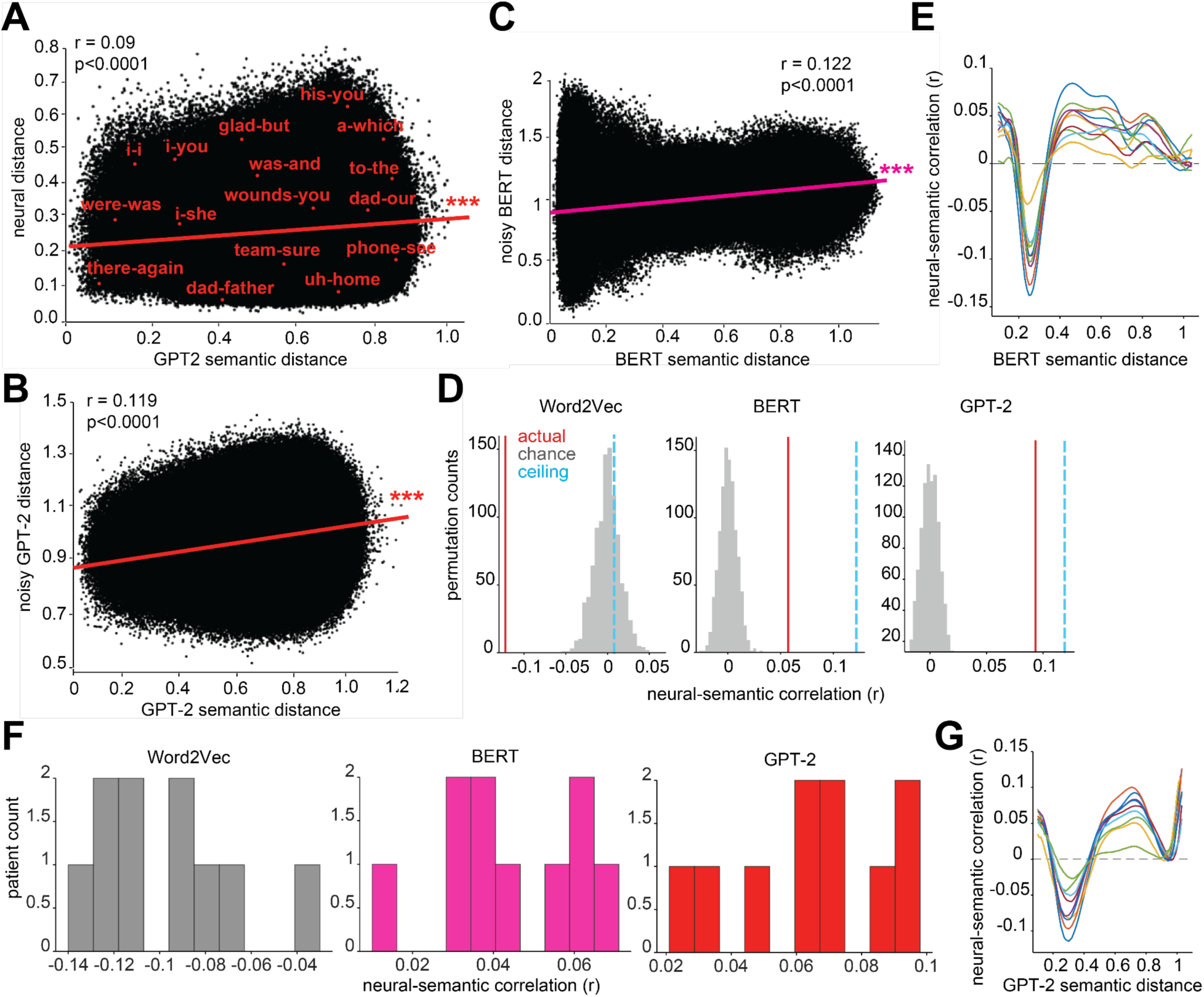
The relationship between neural and semantic distance across language models, noise models, and individual patients. A, Scatter plot illustrating the correlation between neural and GPT-2 layer 37 semantic embedding distances between all 26,978,185 word pairs. Pink line: regression line. Each dot represents one randomly selected word pair; magenta text indicates example pairs. **B,** Scatter plot illustrating the correlation between GPT-2 layer 37 semantic embedding distances and GPT-2 embedding distances that were transformed with neural noise**. C,** the same as in B, but with BERT embeddings from layer 13. **D,** Plots for each language model show the distribution of correlations between neural and semantic distances obtained from shuffled neural activity (gray), the correlation from the real neural and semantic data (red), and the neural noise ceiling (blue dashed line), computed as the correlation between the semantic embeddings and their noise-injected versions. For BERT and GPT-2, the real neural correlation is well above chance and falls below - but close to - the noise ceiling, indicating that the models capture more than half of the explainable neural-semantic structure. In contrast, Word2Vec shows a negative correlation and a modest noise ceiling, consistent with its poor alignment with neural structure. **E,** For each patient (colored line), the semantic-neural correlation coefficient computed at increasing BERT semantic distances (binwidth 0.2, step size 0.01). While this line is generally positive in value, at especially short semantic distances (low x-values), it turns negative, reflecting contrastive coding in each patient. **F,** For each language model, histograms show the neural and semantic distance correlation using neural embeddings from each patient separately. **G,** Contrastive coding for each patient, as computed in panel E but using GPT-2 embeddings.

**Extended Data Fig. 8.**
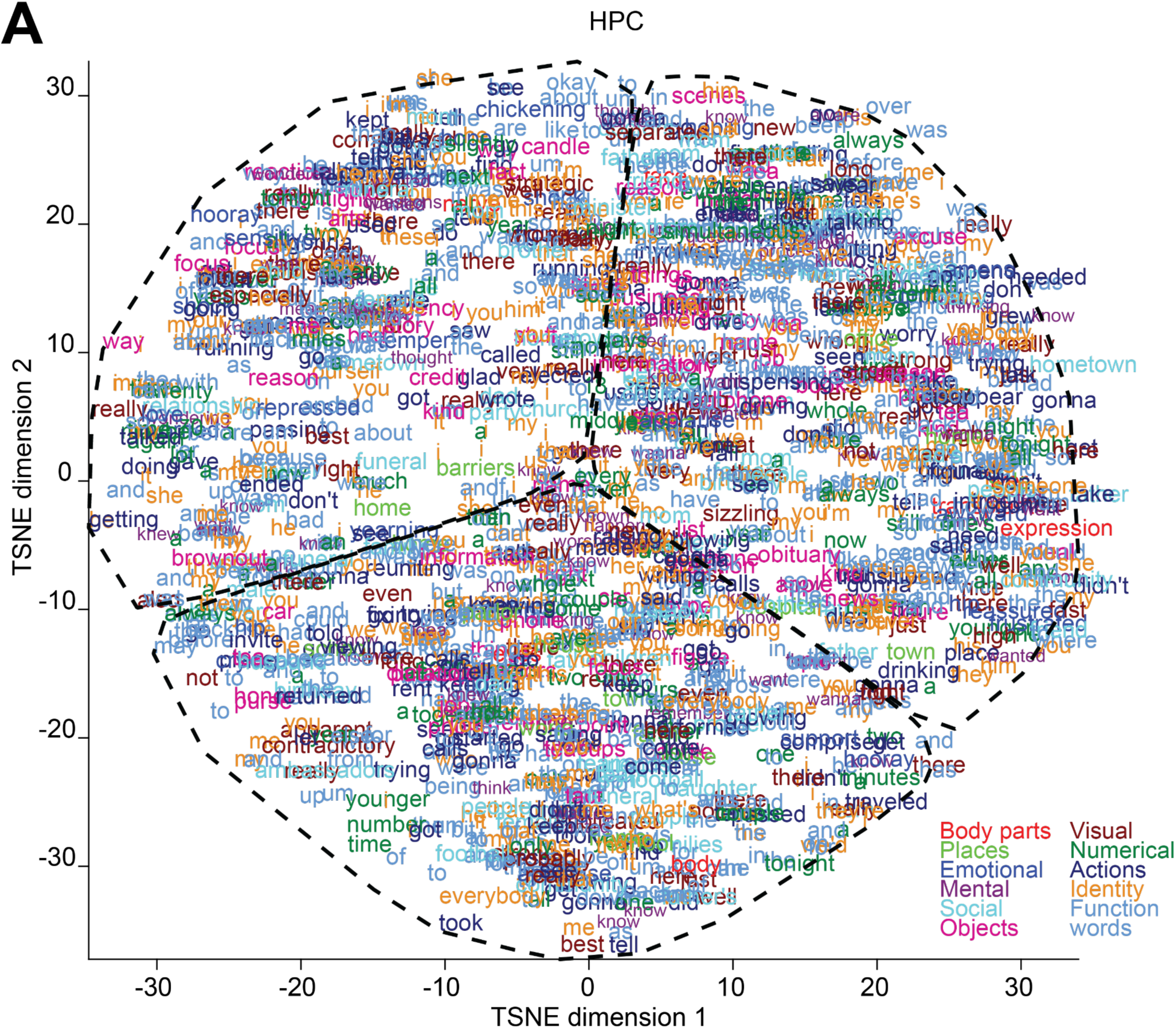
Semantic categories from neural embeddings. A, Words plotted in their dimensionality reduced neural embedding space and colored according to their semantic category defined in Word2Vec. Words are grouped in 3 distinct categories defined from silhouette score and k-means clustering of neural embeddings. Black dashed lines outline the identified categories. 1,808 words out of 7,346 are plotted here for visualization.

**Extended Data Fig. 9.**
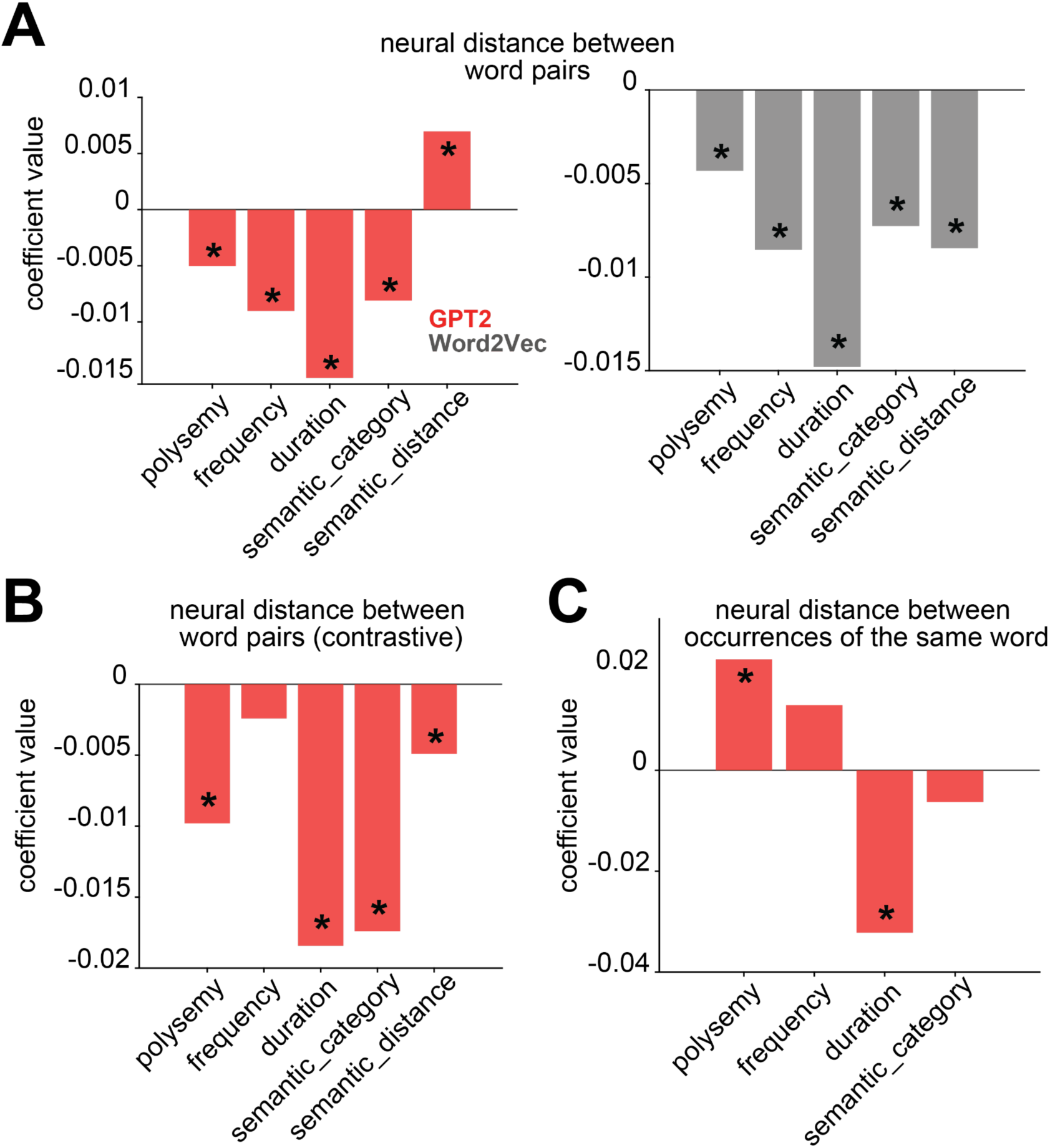
The relationship between linguistic features and neural distance. A, Weights from a ridge regression model that predicted neural distance between all word pairs from the difference in polysemy, word frequency, word duration, and semantic category for each word pair, and the GPT-2 layer 37 (left) or Word2Vec (right) semantic distance between them. Asterisks represent significant predictors. **B,** The same as in A-left but restricted to word pairs with semantic distances within the contrastive coding range, 0.14 - 0.4. **C**, Weights from a ridge regression model that predicted, for every word, the median neural distance between all occurrences of that word from that word’s polysemy (computed with GPT-2 layer 37 embeddings), frequency, duration, and semantic category. Asterisks represent significant predictors.

**Extended Data Fig. 10.**
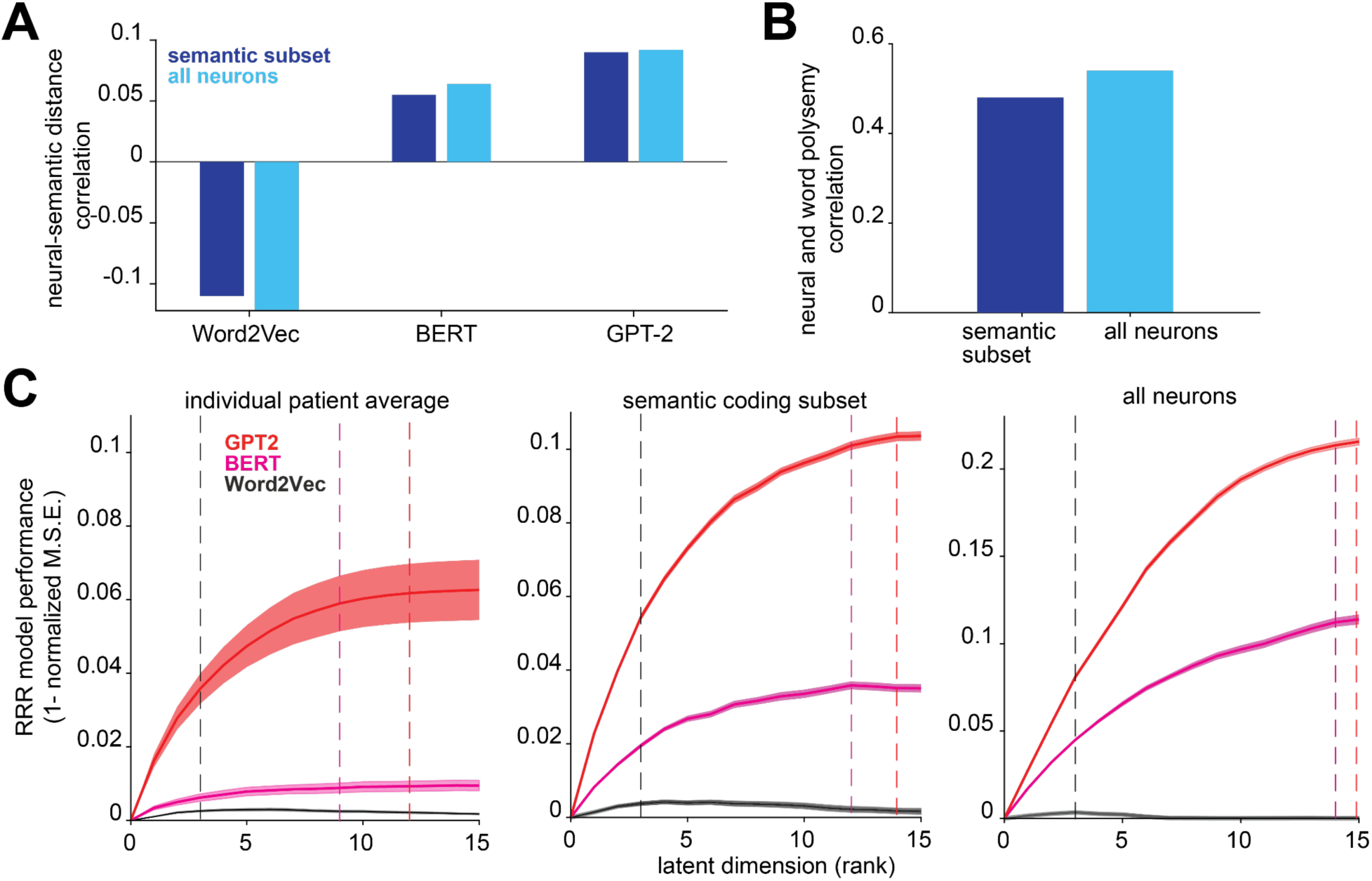
**Comparison between semantic coding using the full population and a semantically selective subset of neurons**. **A,** For semantic embeddings from Word2Vec, BERT layer 13, and GPT-2 layer 37, the Pearson correlation between neural and semantic distance using the the full population of neurons (n = 356) or a subset of neurons whose residual activity was significantly predicted by semantic embeddings (n = 92). All r-values are significant, P < 0.001. **B,** The Pearson correlation between word and neural polysemy, where neural polysemy was computed using a subset of semantically selective neurons or the full population. All r-values are significant, P < 0.001. **C,** Reduced rank regression performance for predicting different language model embeddings from neural population activity using data from each patient individually (n ∼ 30 neurons per patient), the subset of semantically selective neurons (n = 92) or the full population (n = 356). All regression models show the same trends in performance and optimal rank across the three language models.

